# Biochar Rewires Root Exudates and the Rhizosphere Microbiome and Its Functionality

**DOI:** 10.64898/2025.12.25.696513

**Authors:** Hanyue Yang, Amina Furrkukh Mughal, Yaqi You

**Affiliations:** Department of Environmental Resources Engineering, SUNY College of Environmental Science and Forestry; Graduate Program in Environmental Science, SUNY College of Environmental Science and Forestry

**Author notes:** Correspondence: Yaqi You 402 Baker Lab, 1 Forestry Dr, Syracuse, NY 13210.

**Keywords:** biochar, root exudates, secondary metabolites, rhizosphere microbiome, plant growth promoting rhizobacteria, soil metabolome, nitrogen cycling, methane cycling

## Abstract

Biochar amendment is beneficial to the soil-plant system; yet the underlying mechanism is not fully understood. Integrating microfluidics and multi-omics and using wheat as a model plant, we found biochar induces differential root exudates, especially complex molecules like secondary metabolites and signaling cues, and rewires the rhizosphere microbiome, evoking a plant-beneficial rhizosphere microbiome centered by diverse plant growth promoting rhizobacteria (PGPR) known to participate in nitrogen cycling, organic matter turnover, nutrient acquisition, phytohormone metabolism, and stress resistance. Network analysis and machine learning identified high co-occurrence between specific root exudates and PGPR. qPCR, 16S rRNA gene sequencing, and untargeted soil metabolomics suggested the restructured rhizosphere microbiome may have shifted nitrogen and methane metabolism leading to reduced potential of N2O and methane emissions, and may participate in the metabolism of signaling cues and redox reactions. This study provides new insights into biochar’s profound impact on the soil-plant system and highlights the potential of engineering the rhizosphere through reshaping root-microbe interactions.

## 1. INTRODUCTION

Sustainable soil management is vital for safeguarding crop production, environmental health, and human well-being.^1^ Biochar has been widely recognized as a multifunctional soil enhancer, capable of improving soil health, boosting crop productivity and stress resistance, reducing nutrient leaching, and mitigating greenhouse gas (GHG) emissions.^2–4^ These benefits are often attributed to biochar-induced changes in soil physicochemical properties, such as increased nutrient availability, enhanced water-holding capacity, and reduced acidity or toxic substances.^2^ Emerging evidence including our research,^5^ however, started to unveil more profound influences of biochar on the soil-plant system.

Soil is a dynamic living system, comprising diverse and complex microbial communities that drive numerous biogeochemical reactions.^6^ Increasing evidence suggests that biochar amendment can alter microbial abundance, diversity, community structure, and functional compositions, as synthesized in our meta-analysis.^3,7^ However, few studies have investigated biochar’s effects on the rhizosphere−a narrow, dynamic zone that serves as a critical interface between plant roots and soil microorganisms and a key component of the plant holobiont.^3,8,9^ Our recent study found that biochar can modulate root endometabolites and rhizosphere microbiome.^5^ But not all root endometabolites are secreted into the soil; root exudates that are actively secreted directly influence rhizosphere chemistry and thus require further investigation.

Plants use root exudates to recruit plant-growth promoting rhizobacteria (PGPR) and thereby gain growth advantages such as enhanced resistance to diseases and stresses.^8,9^ Associations between biochar-induced root exudates and PGPR enrichment is still unknown. Further, nitrogen (N) is a key limiting nutrient for plant productivity in most terrestrial ecosystems, especially croplands.^10,11^ While biochar is reported to affect microbially mediated N cycling, including N fixation, nitrification, and denitrification, in bulk soil,^3,12^ its effect on rhizosphere N cycling is not well understood. Similarly, biochar’s effect on rhizosphere methane cycling is poorly understood. Specifically, the role of biochar-induced root exudates therein is understudied. Unleashing biochar’s potential in sustainable crop production requires a mechanistic understanding of biochar’s modulating effects on root exudation and the rhizosphere microbiome.

This study aimed to use wheat (*Triticum aestivum L.*), a fertilizer-intensive crop, as a model plant to elucidate biochar-induced chemical and biological shifts in the rhizosphere, with a focus on molecular mechanisms of biochar-induced rhizosphere reprogramming. Three specific objectives were: (1) identify biochar-induced root exudates, (2) characterize biochar-induced, root exudate-mediated rhizosphere microbiome restructuring, and (3) determine biochar’s modulation on rhizosphere biogeochemistry, especially N and methane cycling. To that end, we conducted controlled experiments using laboratory-produced, well-characterized wheat biochar and a microfluidics device called EcoFAB (ecosystem fabrication), a modular growth system developed for studying the rhizosphere throughout plant development stages.^13^ We analyzed root exudates and the soil metabolome using high-resolution mass spectrum (HRMS)-based untargeted metabolomics, the rhizosphere microbiome and its potential function using next-generation sequencing and real-time quantitative PCR (qPCR), and root exudate-microbe associations using a combination of network analysis and machine learning. Our work provides new mechanistic insights into biochar’s multifaceted roles in reprogramming the rhizosphere and the potential of engineering the rhizosphere through reshaping root-microbe interactions.

## 2. MATERIALS AND METHODS

### 2.1. Biochar production and characterization, and EcoFAB fabrication

Biochar was produced by slow pyrolysis of wheat straw (350 °C for 30 min) in a laboratory reactor, the same as our prior work.^5,14^ We chose the pyrolysis temperature due to the fact that biochar produced at lower pyrolysis temperatures tends to promote soil enzyme activity more than biochar produced at higher pyrolysis temperatures.^3^ Biochar was characterized using conventional (e.g., pH, N species) and advanced approaches (e.g., Fourier-transform infrared spectroscopy, X-ray diffraction) (details in the Supporting Information (SI)).

We used EcoFAB devices, a fabricated ecosystem, for wheat plant cultivation, rhizosphere observation, and sample collection.^13,15^ This setting enables continuous investigation of the rhizosphere under highly controlled conditions.^16^ We downloaded the original design files (https://eco-fab.org/device-design/) and customized the design to accommodate wheat seedlings. Acrylonitrile butadiene styrene (ABS) mold parts were fabricated using a MakerBot Replicator 2X 3D printer, acrylic mold parts were fabricated on a lathe machine, and the assembled mold was used to cast polydimethylsiloxane (PDMS) layers (**Fig 1A**). Our prototype EcoFAB device consisted of a PDMS oval chamber (97.6 × 64 × 3.2 mm) adhering to a large rectangular coverslip (102 × 76 × 1.2 mm) (Ted Pella) in an airtight way (**Fig 1A**). Each device was enclosed in a transparent cover box to maintain a sterile environment for plant experiment (**Fig 1B**). Before experiments, the EcoFAB chamber was rinsed with sterile Milli-Q water; the device surface and the cover box were sterilized using 70% ethanol for 30 minutes and then 100% ethanol for 5 min. More details about EcoFAB fabrication and device assembly are in the SI.

**Fig 1.**
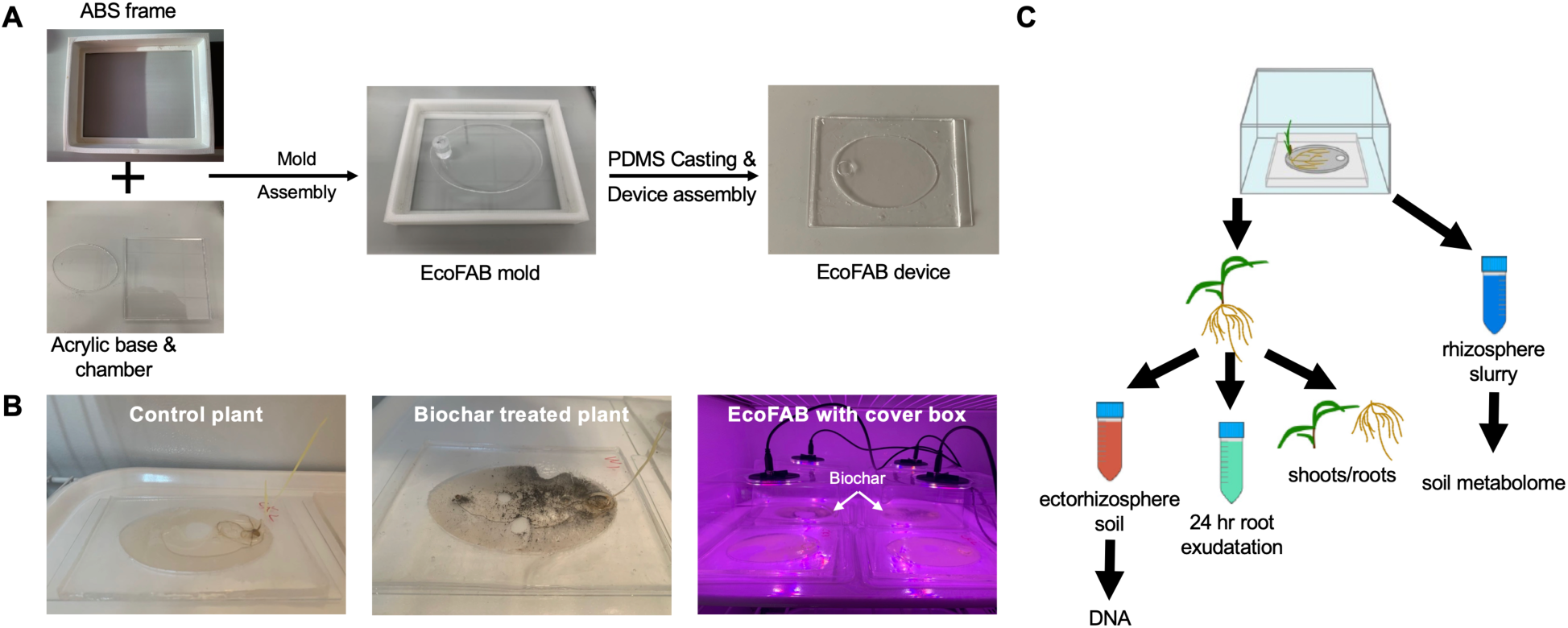
(A) Fabrication of a prototype EcoFAB device. (B) EcoFAB experiment under a sterile and highly controlled environment. The EcoFAB chamber hosted the root system while the plant shoot was covered by a sterile cover box equipped with a LED (full spectrum) control and a humidity monitor. Each EcoFAB device represents a biological replicate, and an individual wheat seedling was cultivated with or without 0.25% (w/w) wheat biochar. (C) Sequential sample collections: (1) rhizosphere soil slurry for the soil metabolome, (2) ectorhizosphere soil for DNA extraction, and (3) 24 hour hydroponic secretion for root exudates after ectorhizosphere soil removal. The schematic of EcoFAB experiment is adapted from Sasse et al. (2019).^16^

### 2.2. Plant experiment and sample collection

We applied wheat biochar at a rate of 0.25% (w/w) based on cumulative evidence that lower rates promote plant growth while higher rates inhibit growth and our prior observation that wheat biochar at 0.25% rate yields unique rhizosphere alterations.^5^ To prepare soil slurries, 100 g of agricultural soil (properties in **Table S1**) was added to 1 L sterile Milli-Q water, with or without biochar. Soil slurries were shaken for 16 hours to reach thorough mixing, and slowly injected into each EcoFAB chamber with a sterile syringe. Wheat seeds were germinated and seedlings of similar sizes were transferred aseptically into sterilized EcoFAB devices containing soil slurries with or without biochar (one plant per device; four biological replicates for each treatment) (**Fig 1B**). Plants were cultivated under full spectrum LED light simulating natural sunlight (12 hour light, 12 hour dark) and controlled humidity (80%) at 25 °C for 21 days, and the development of root systems was monitored (**Fig 1B**). Milli-Q water was added to EcoFAB devices every three days to maintain humidity. To ensure reproducibility, the experiment was conducted twice.

After 21 days, individual EcoFAB devices were dissembled, and remaining soil slurries and plants were collected (sequential sample collections shown in **Fig 1C**). Soil slurries were immediately stored at -80°C until metabolite extraction, while plants were immediately processed to collect ectorhizosphere soils like in our prior studies.^5,17^ After ectorhizosphere soil removal, individual plants with clean root surfaces were transferred to sterile 15 mL centrifuge tubes containing 15 mL Milli-Q water to allow for exudation for 24 hours like in our prior study.^18^ An aliquot of the collected exudated solution (1.5 mL) was transferred to a 2.0 mL sterile microcentrifuge tube, dried using a vacuum concentrator (Savant SpeedVac SPD300DDA, Thermo Scientific), and 1.5 mL of methanol solution (80% v/v in Milli-Q water) was added to the tube, followed by vortexing for 15 minutes and sonication bath for 5 minutes at room temperature. The resulting extract was filtered through a 0.22 µm PVDF filter (Millipore) and stored at -80 °C until metabolomics analysis. Similar procedures were used to extract soil exometabolites (detailed in the SI).

### 2.3. Untargeted metabolomics using HRMS

Root exudates and soil exometabolites were analyzed using liquid chromatography-mass spectrometry (LC-MS) performed on a Vanquish LC system in conjunction with a Q Exactive HF Orbitrap MS coupled with an electrospray ionization (ESI) source (Thermo Fisher). The MS has a resolving power of up to 240,000 FWHM (Full Width at Half Maximum) at 500 m/z and a mass range of 20-1200 m/z. Extract aliquots of 10 µL were injected into a Hypersil GOLD C18 column (1.9 µm, 2.1×150 mm) (Thermo Fisher) operated at 30 °C. LC mobile phases consisted of solvent A (0.1% formic acid in Milli-Q water) and solvent B (0.1% formic acid in LC/MS grade acetonitrile), and the flow rate was maintained at 300 µL/min. MS was operated in both positive and negative ion mode (ESI+ and ESI-). More LC-MS details are in the SI. Spectral data was processed as before.^5^ Raw data was converted to the generic mzXML format and processed using three platforms (MetaboAnalyst 6.0, XCMS, MZmine3).^19–21^ The resulting feature tables were merged without redundancy and further processed in MetaboAnalyst 6.0 for functional analysis. Compounds were converted to the International Union of Pure and Applied Chemistry (IUPAC) names and classified to the “Class” level of the purely structure-based chemical taxonomy (ChemOnt) using ClassyFire.^22^ Significant fold change (FC) was determined by the Mann-Whitney *U* test with Benjamini-Hochberg correction (|log_2_FC| > 1 and *p* < 0.05). Principal coordinates analysis and network analysis are detailed in the SI.

### 2.4. Next generation sequencing, qPCR, and biofilm assay

Metagenomic DNA was extracted from rhizosphere soils using the ZymoBIOMICS DNA/RNA Miniprep Kit following the manufacturer’s protocol (Zymo Research). A PowerLyzer Homogenizer was used to maximize extraction efficiency. The resulting DNA was assessed on a Qubit 4 fluorometer with the 1× dsDNA High Sensitivity Assay Kit (Thermo Fisher) and by electrophoresis. 16S rRNA gene sequencing and data processing (QIIME2, DADA2, phyloseq) were performed as before.^5,23–25^ The “edgeR” package was used to identify differential taxa between samples. PICRUSt2 was used to infer metagenomic functional contents; PICRUSt2 predictions were subjected to differential abundance analysis using the R package “ggpicrust2”.^26,27^ Microbial community growth rate, i.e., median predicted minimum doubling time (mPMDT), was predicted using the R package “gRodon”.^28,29^ Microbial network analysis was constructed using SparCC.^30^ Community assembly mechanisms were assessed using the null model approach. We calculated the β-nearest taxon index (βNTI) using the R package “iCAMP” to estimate the relative importance of deterministic versus stochastic assembly processes,^31^ and calculated the Raup-Crick metric using Bray-Curtis dissimilarities (RC_BC_) to estimate the relative importance of selection, drift, dispersal limitation, and homogenizing dispersion.^32^ More details on sequencing and data processing are in the SI.

While PICRUSt2 inference provides a more comprehensive snapshot of a community’s potential function, inference accuracy varies. Therefore, we used qPCR to quantify the abundance of eubacterial 16S rRNA gene (proxy of total bacteria) and key genes involved in nitrogen (*nifH*, *nxrA*, *nirK*, *nirS*, *nosZ* Clade I) and methane (*mcrA*, *pmoA*) cycling using established primers (**Table S2**). qPCR was performed on a CFX Duet Real-Time PCR System (BioRad) with SsoAdvanced universal SYBR green supermix (Bio-Rad), 2 μL of DNA template (2-fold diluted to mitigate soil matrix inhibition effect), and primers (0.48 μM each) in a 25 μL reaction volume (details in the SI).

To test whether biochar-induced root exudates promote biofilm formation compared to root exudates from control plants, we conducted a biofilm assay using a model soil bacterium *Pseudomonas putida* KT2440. The results were normalized against dissolved organic carbon (DOC) content in root exudates to exclude the effect of varying carbon content (details in the SI).

### 2.5. Multi-omics integration

We used two approaches to identify associations between root exudates and rhizosphere microbes: (1) OmicsNet 2.0 which utilizes multiple databases to create multi-omics networks, and (2) mmvec (microbe-metabolite vectors), a neural network approach that predicts an entire metabolite profile from a single microbe sequence.^33,34^ OmicsNet 2.0 calculates metabolite-microbe correlations based on microbial metabolic capacity and pertinent proteins that are predicted using high-quality genome-scale metabolic models. The mmvec approach estimates the conditional probability of observing a metabolite given that a microbe is present to identify the most likely metabolite-microbe interactions. Compared to approaches based on correlation (e.g., Spearman’s) or network (e.g., SparCC), mmvec can identify previously undiscovered patterns and is less prone to false discovery (positive or negative) as it provides consistency between absolute and relative abundances of compositional data. However, mmvec does not calculate the statistical significance or confidence interval of each metabolite-microbe pair. Here we combined the two approaches to maximize the discovery of true metabolite-microbe interactions. Similar approaches were used to identify and visualize associations between rhizosphere microbes and soil exometabolites. More details are in the SI.

### 2.6. General statistics and data visualization

Bioinformatics, statistics, and data visualization were performed in R (4.2.1) and Python (3.7) with various R packages and Python modules. Significant differences were determined using one-way analysis of variance (ANOVA) or Welch’s *t* test, nonparametric statistics (Wilcoxon or Kruskal-Wallis), and permutational multivariate analysis of variance (PERMANOVA), depending on the variable. Network visualization used Gephi (0.10.1) and Cytoscape (3.9.1). More details are in the SI.

## 3. RESULTS AND DISCUSSION

### 3.1. Biochar significantly promotes plant growth

SEM indicated the wheat biochar used here had a porous morphology (**Fig 2A**). FTIR spectra showed the biochar contained aromatic compounds (particularly those with benzene rings and quinone groups), C–O bonds from esters or ethers, nitriles, silicates and/or carbonates, and possibly C–N or P–O (**Fig 2B; Fig S2**) (more in the SI).^35,36^ EDS analysis revealed the surface of the biochar predominantly contained C (62.36%), O (24.52%), Si (6.13%), K (3.06%), and Ca (1.04%) (averaged atomic%, n = 29) (**Fig 2C**). Consistent with the EDS results, XRD spectra indicated the presence of amorphous C, graphite, and various mineral crystals containing Si, K, and Ca (**Fig 2D**) (more in the SI).^36,37^ These results reflect the biochar’s feedstock (wheat straw) and the production condition (slow pyrolysis at 350 °C).

**Fig 2.**
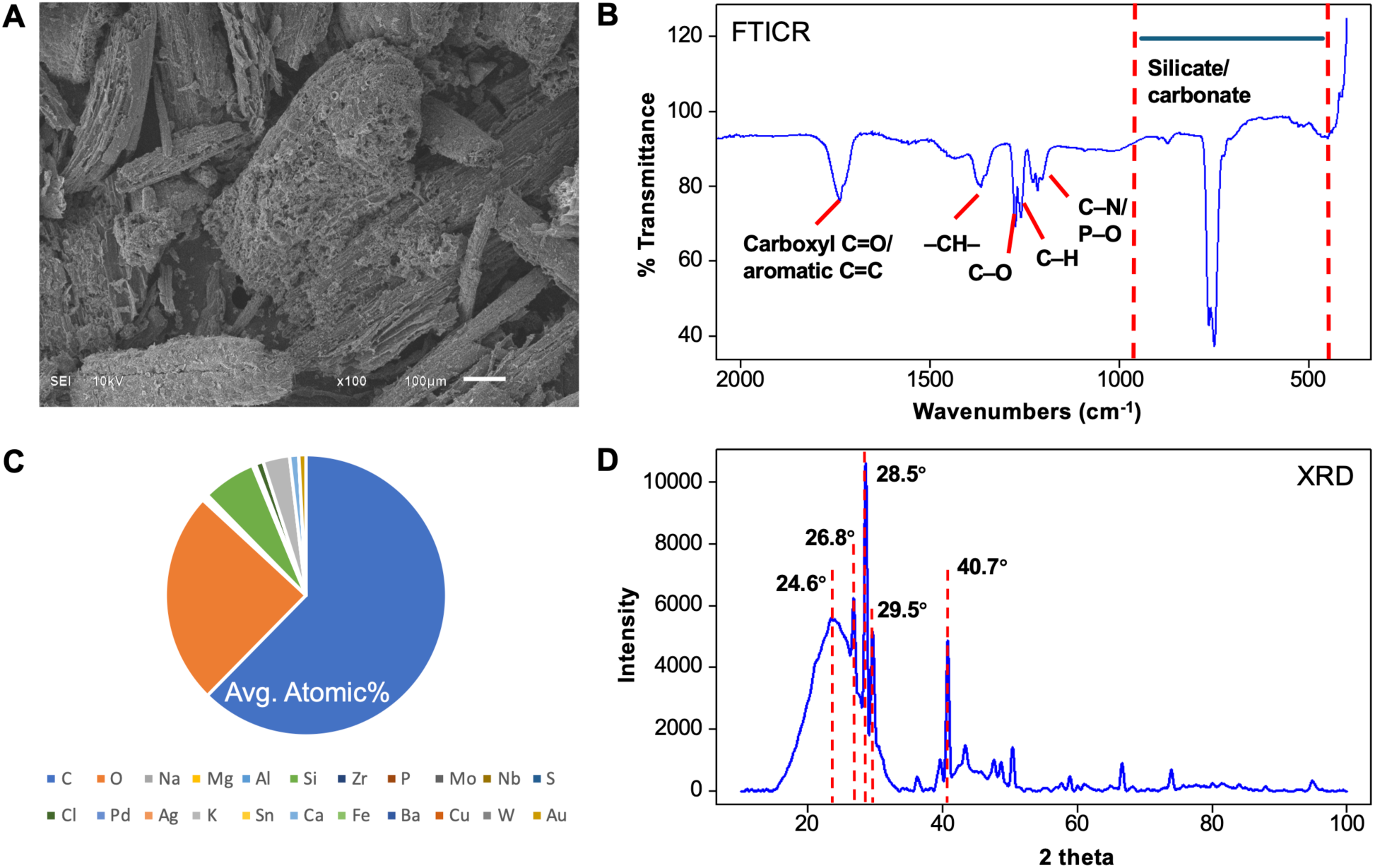
(A) Representative SEM image of the wheat straw biochar used in this study. (B) FTIR spectrum (400-2000 cm^-1^) with annotations of characteristic peaks (see **Fig S2** for full spectrum image). (C) EDS-derived elemental distribution on biochar surface (atomic percentages averaged from 29 subsamples). (D) XRD spectrum with characteristic diffraction angles.

In the 21-day EcoFAB experiment, application of 0.25% (w/w) wheat biochar yielded 1.59 times more fresh weight of shoots (Welch *t* test: *p* < 0.001) (**Fig S3**). Fresh weight of roots had no significant difference (Welch *t* test: *p* = 0.413), but biochar-treated plants had more lateral roots (**Fig S3**). These results are consistent with our prior 15-day pot study and suggest biochar’s plant growth promotion effect may be scalable.^5^

### 3.2. Biochar induces differential root exudation

Biochar induced differential root exudation (**Table S3**), systematically regulating root metabolism, as evidenced by increased average degree and centrality of root exudate co-occurrence network (**Fig 3A**). Specifically, biochar significantly regulates carbohydrate metabolism, glycan biosynthesis, the metabolism of terpenoids and polyketides, and nucleotide metabolism, all of which are highly involved in plant growth and development, signaling, and stress defense.^2,3^ Among the 312 compounds detected across samples, six were only identified in biochar-induced root exudates, including two flavanonols, one flavanone and its direct precursor, a phenolic ester, and dUMP (**Table S3**). Further, biochar significantly affected 39 compounds from 14 ChemOnt classes (|Log_2_FC| > 1, *p-adjusted* < 0.05): 10 organooxygen compounds, 6 flavonoids, 3 prenol lipids, 3 fatty acyls, 2 from each of four classes (organonitrogen compounds, steroids and steroid derivatives, imidazopyrimidines, carboxylic acids and derivatives), 1 from each of five classes (5’-deoxyribonucleosides, pyrimidine nucleotides, furanoid lignans, linear 1,3-diarylpropanoids, organic phosphoric acids and derivatives), and 4 unclassed chemicals (**Fig 3B**). Two compounds having regulatory roles in plant defense were upregulated, including (22S)-22-hydroxycampest-4-en-3-one (a direct precursor of brassinosteroids (BRs)) (2.6 fold) and 2,4-dihydroxy-1,4-benzoxazin-3-one-glucoside (DIBOA-Glc, a benzoxazinoid (BX) compound) (5.2 fold). BRs are phytohormones indispensable for root growth and BR biosynthesis is largely restricted to the root elongation zone.^38^ DIBOA-Glc is the glycosylation product of DIBOA, the most abundant BX in cereal crops, and the precursor of defense-active compound DIMBOA-Glc.^39^ BX glucosides are stored in vacuoles and released over time to the rhizosphere where their derivatives can suppress weeds and hostile microbes while enriching beneficial microbes.^40,41^ These results indicate that biochar may promote plant growth by upregulating root exudates that have antimicrobial and antioxidant activities. Further, BRs and BXs are secondary metabolites known for their signaling functions, and increasing evidence reveals they also mediate plant-microbe interactions such that shifts in their biosynthesis can reshape the rhizosphere microbiome.^40–44^ Here, by upregulating biosynthesis of plant signaling metabolites, biochar may modulate the rhizosphere microbiome, which indeed occurred as discussed later. Thirty-six root exudates were downregulated, including six flavonoids. Flavonoids are essential in plant fitness and rhizosphere microbiome assembly.^45^ Downregulation of flavonoid biosynthesis could reflect either reduced stress due to scavenging of reactive oxygen species or BR-induced inhibition of flavonoid biosynthesis.^46^ It may also reflect shifted flavonoid biosynthesis as several flavonoids, including 2 flavanonols (dihydrokaempferol, (+)-2,3-dihydrofisetin), one flavanone and its direct precursor (eriodictyol and eriodictyol chalcone), and a phenolic ester (4-coumaroylshikimate) were only detected in biochar-treated root exudates. Overall, our results provide new chemical insights into widely observed biochar benefits on crop productivity.^47^

**Fig 3.**
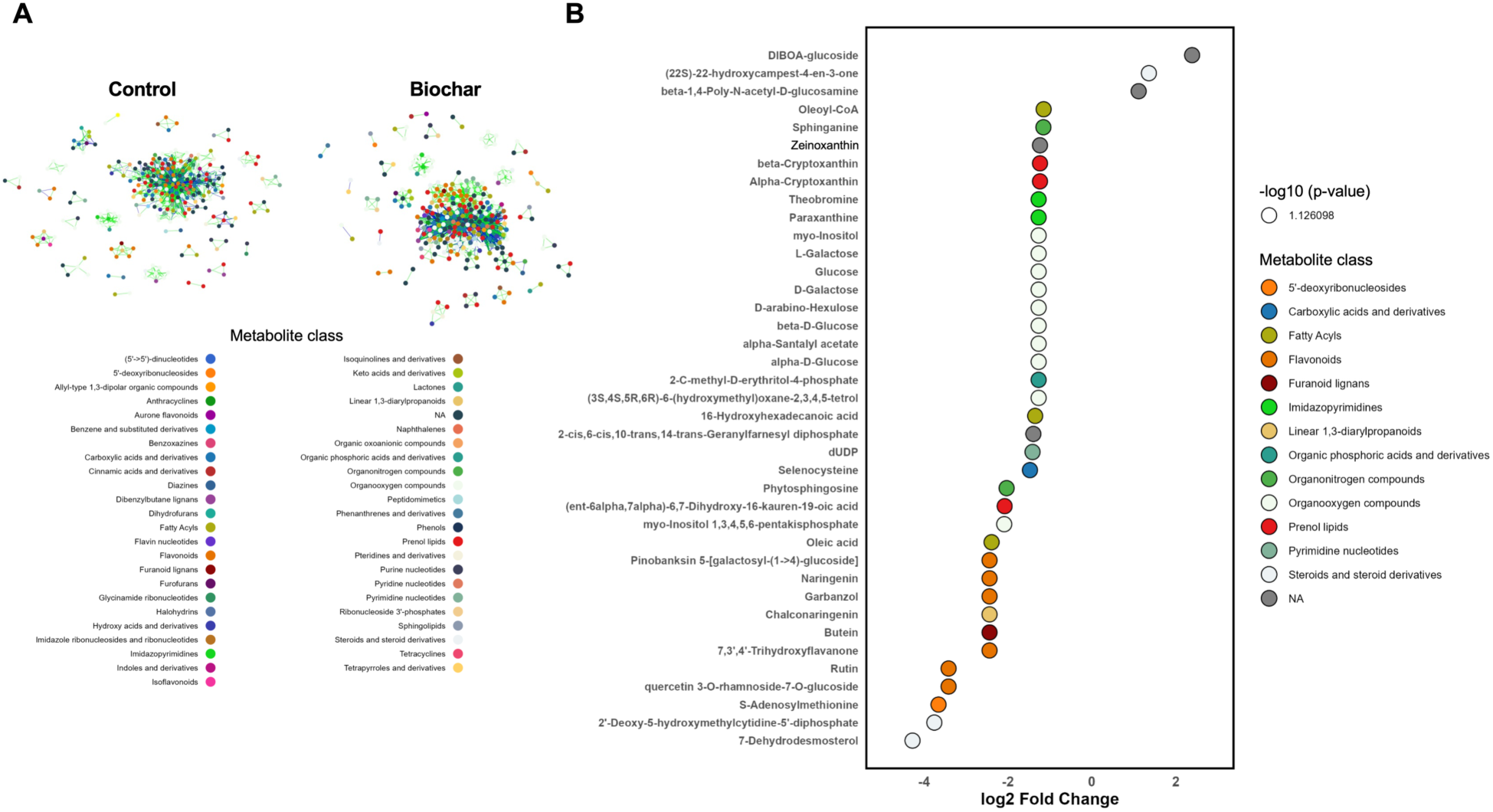
(A) Co-occurrence network of root exudates of the control and biochar-treated plants (*r* > 0.9, *p-adjusted* < 0.01). Dots indicate individual root exudates colored based on ChemOnt classes. (B) 39 most discriminative root exudates from biochar treatment (|Log_2_FC| > 1, *p-adjusted* < 0.05). Compounds are grouped into ChemOnt classes.

A majority (81.4%) of the 312 exudate compounds were previously detected inside of root tissues (**Fig S5**; the SI text).^5^ But 58 exudate compounds were not previously detected within roots, including 8 fatty acyls, 6 organooxygen compounds, 5 pyrimidine nucleotides, 5 purine nucleotides, 4 carboxylic acids and derivatives, 3 flavonoids, 1 isoflavonoid, 3 prenol lipids, among others, such as DIBOA-Glc and gibberellin A29-catabolite (GA29-catabolite, a deactivated catabolite from GA20) (**Fig S5; Table S4**). These 58 actively exudated compounds have critical roles in the rhizosphere: Fatty acyls and prenol lipids, organooxygen compounds, carboxylic acids and derivatives, and flavonoids and isoflavonoids are involved in rhizosphere signaling, able to mediate plant root-microbe interactions.^9,45,48^ Among the detected fatty acyls, linolenic acids are plant defense signaling molecules;^49^ 2’-deoxymugineic acid is an iron (III) phytosiderophore, influencing rhizosphere microbiome composition through iron competition.^8,50^ Five of the six detected organooxygen compounds were inositol phosphates, a major pool of organic phosphorus in soil and signaling molecules mediating microbial root colonization.^51^ These results, along with our prior pot study,^5^ suggest biochar-induced metabolic alterations in roots and subsequent exudation of exudates that may modulate plant-microbe interactions.

### 3.3. Rhizosphere microbiome restructuring and its association with biochar-induced root exudates

#### 3.3.1. Biochar rewires the rhizosphere microbiome and microbial interactions

In the native wheat rhizosphere, the top 10 phyla and top 40 genera comprised 60% and 35% of all prokaryotes, respectively, reflecting the core rhizosphere microbiome of plant, especially cereals.^8^ Biochar had limited influence on rhizosphere microbial alpha diversity but considerably altered microbiome composition (PERMANOVA test: *p* = 0.001 for round × treatment) (**Fig 4A-B**), promoting 12 genera while suppressing 9 genera (**Fig S6-S7; Table S5**) (more in the SI).

**Fig 4.**
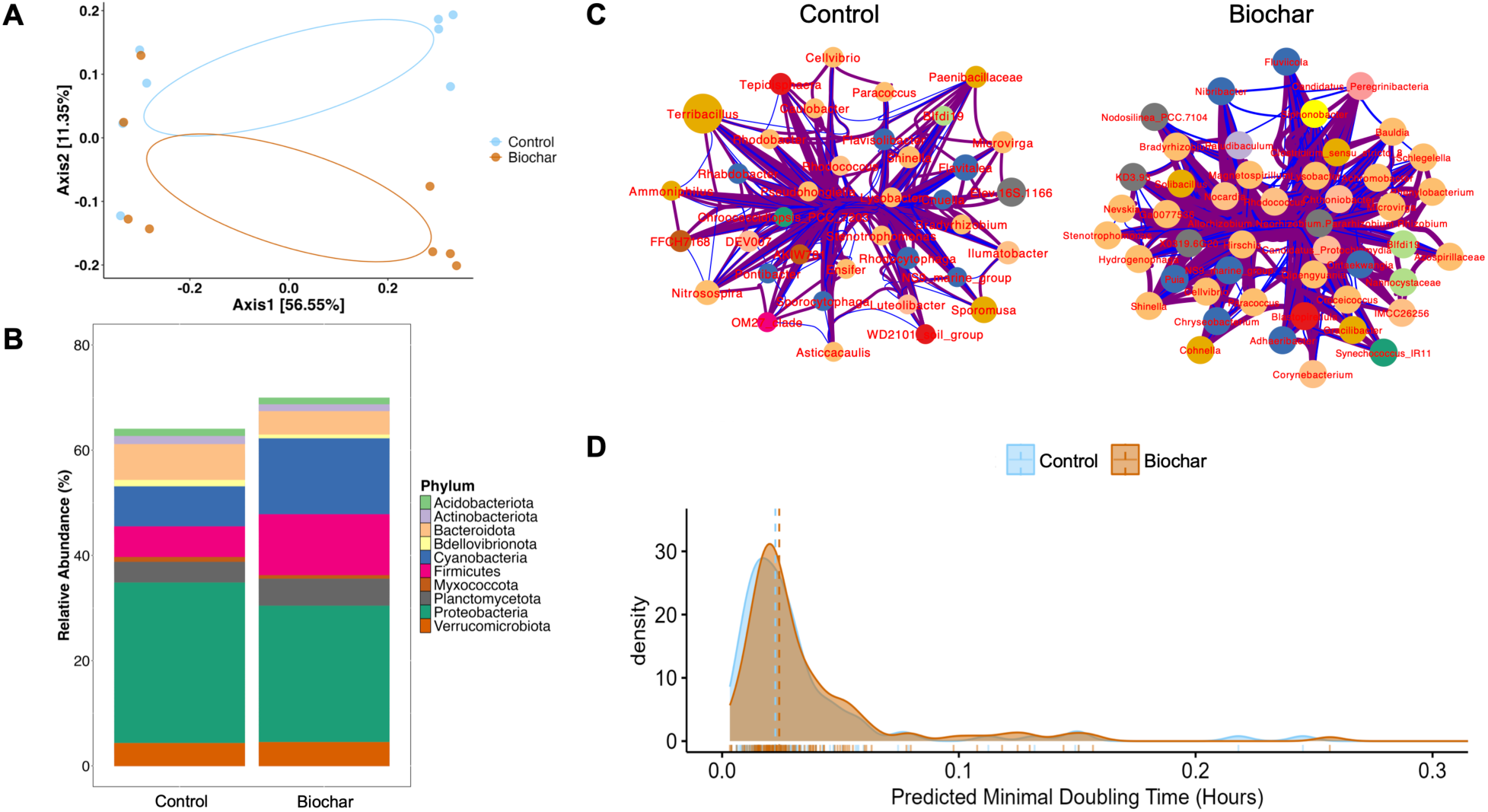
(A) PCoA of the native and biochar-treated rhizosphere microbiomes (PERMANOVA test: *p* < 0.05 for round x treatment). (B) Relative abundance of the top 10 microbial phyla. (C) Hub networks of keystone genera in the native and biochar-treated rhizosphere (*p* < 0.05, 1000 bootstrapping). Node colors and sizes indicate phyla and betweenness values. Purple and blue edges indicate positive and negative correlations. (D) Biochar-rewired rhizosphere microbiome had slightly slower growth rate (i.e., larger mPMDT) than the native rhizosphere microbiome based on gRodon^28,29^ prediction using enriched taxa (DESeq2: |Log_2_FC| > 1, *p-adjusted* < 0.05).

Remarkably, biochar enhanced microbial interactions, evidenced by greater microbial network modularity and centrality and a large number of newly emerged keystone genera (60 vs. 53 in the treated vs. native rhizosphere) (**Fig 4C; Fig S8; Table S6-S8**). Twelve common keystone bacterial genera were observed in the native and treated rhizosphere, including nine PGPR from Actinobacteria, Alphaproteobacteria and Gammaproteobacteria: *Bradyrhizobium* and *Microvirga* are N-fixing bacteria;^52^ *Paracoccus* can metabolize nitrogen, nitrate, and ammonia;^53^ *Lysobacter* and *Stenotrophomonas* can suppress soil-borne pathogens by producing lytic or antimicrobial substances;^54,55^ *Shinella* can degrade polycyclic aromatic hydrocarbons;^56^ *Cellvibrio* has an extensive repertoire of carbohydrate-active enzymes for degrading polysaccharides (cellulose, dextran, xylan, chitin, and starch), thus enhancing organic matter turnover;^57^ *Rhodococcus* contributes to N cycling and influences plant stress response and growth by metabolizing the phytohormone abscisic acid (ABA) and producing the phytohormone indole-3-acetic acid (IAA);^58,59^ *Solirubrobacter* can produce IAA and improve nutrient bioavailability through assimilatory reduction of sulphate and utilizing a wide range of carbohydrates (more in the SI).^60^

Besides these common keystone genera, the native rhizosphere microbiome had classic rhizosphere keystone genera, such as those able to degrade plant cell wall polysaccharides (detailed in the SI), whereas the biochar-treated rhizosphere had additional diverse PGPR as keystone genera (**Fig 4C**). These included *Chryseobacterium* (with greatest betweenness and closeness centrality) that can enhance plant nutrient acquisition, degrade aromatic compounds, produce antioxidant enzymes, synthesize IAA, and lower plant stress levels by reducing the immediate precursor of the stress phytohormone ethylene using 1-aminocyclopropane-1-carboxylate (ACC) deaminase.^9,61^ Another example was methylotrophic *Methylobacterium*/*Methylorubrum* (connector hub) that can synthesize developmental phytohormones (auxins, gibberellins, cytokinins) and ACC deaminase and fix nitrogen.^9,62^ Other newly emerged keystone PGPR genera in the biochar-treated rhizosphere included N-fixers *Allorhizobium*-*Neorhizobium*-*Pararhizobium*-*Rhizobium* and *Azospirillaceae*,^40^ denitrifiers *Magnetospirillum*,^63^ IAA-producing *Nocardia*,^64^ Planctomycetes *OM190* (module hub) able to produce diverse secondary metabolites including antimicrobial compounds,^65^ and *Solibacillus* that can produce IAA and plant singling molecule hydrogen cyanide and secrete siderophores to solubilize or chelate minerals.^66^

These results suggest a potential microbial mechanism for the widely reported but poorly understood plant-promoting effect of biochar: It elicits diverse PGPR, including nitrogen fixers, denitrifiers, and phytohormone producers, to emerge as keystone taxa. Therefore, the biochar-treated rhizosphere microbiome may grant the plant with multiple benefits.^67^ Indeed, biochar-treated plants had higher shoot mass and more lateral roots (Section 3.1), a phenomenon likely attributed to the enrichment of PGPR and the upregulation of BR biosynthesis (Section 3.2). Given the significance of keystone taxa in overall microbiome structure and functioning,^8^ future research would benefit from directly characterizing functions of the keystone microbes in the biochar-treated rhizosphere.

3.3.2. *Rhizosphere microbiome rewiring is associated with biochar-induced root exudates*

Plants use root exudates to recruit and tune the rhizosphere microbiome.^8^ Since biochar induced differential exudation, particularly polyphenols and lipids, we hypothesized that shifts in root exudate chemistry may be associated with rhizosphere microbiome restructuring, selectively enriching or suppressing bacterial taxa, through metabolic coupling of biochar-induced root exudates with microbial substrate-use preferences as suggested by recent evidence from laboratory experiments and genome-informed modeling.^68–70^ In line with this hypothesis, computational prediction showed the biochar-rewired rhizosphere microbial community had a slower growth rate than the native microbiome (**Fig 4D**), likely due to a switch from higher growth and lower substrate utilization efficiency to lower growth and higher substrate utilization efficiency.^68,70^ To validate this genome-informed, community-level prediction, we tested biofilm formation of a model soil bacterium *P. putida* KT2440 under exposure to control or biochar-induced root exudates (**Fig S9A**). While root exudates from control and biochar-treated plants had similar levels of DOC (**Fig S9C**) (*p* = 0.607), biochar-induced root exudates resulted in less *P. putida* biofilm within 48 hours (**Fig S9B**) (*p* = 0.004), consistent with the computational prediction that biochar-induced root exudates favor slow-growing microbes capable of utilizing complex molecules. It should be noted that this shift did not significantly change the total bacteria number. Instead, biochar insignificantly increased the number of rhizosphere bacteria as shown by qPCR results (**Fig S10**). Analysis of community assembly mechanisms found that dispersal limitation dominated rhizosphere community turnover (**Fig S11A-B**), suggesting low dispersal rates in the rhizosphere niche where weak or unstructured selective pressures from substrate utilization enable high compositional turnover.^31^ Together, computational prediction, biofilm assay, and community assembly analysis support our hypothesis that biochar rewired the rhizosphere microbiome through metabolic coupling of biochar-induced root exudates with microbial substrate-use preferences.

Our hypothesis is further supported by the coincidence of biochar-upregulated root exudates like DIBOA-Glc and biochar-enriched bacteria having BXs-enhanced growth (detailed in the SI). This motivated us to explore specific microbe-exudate compound associations. Combining the co-occurrence network (OmicsNet 2.0) and neural network (mmvec) approaches, we identified 12 bacterial genera, including 11 PGPR (*Bacillus*, *Blastococcus*, *Bradyrhizobium*, *Brevundimonas*, *Mesorhizobium*, *Nevskia*, *Paenibacillus*, *Pantoea*, *Pseudomonas*, *Rhodococcus*, *Roseomonas*),^52^ with highest associations with biochar-induced root exudates (mainly plant secondary metabolites) (**Fig 5**). Extremely high co-occurrence probability may manifest when a compound is a preferred substrate to certain bacteria,^40,68–70^ and the root exudate-microbe associations observed here are in line with existing literature.

**Fig 5.**
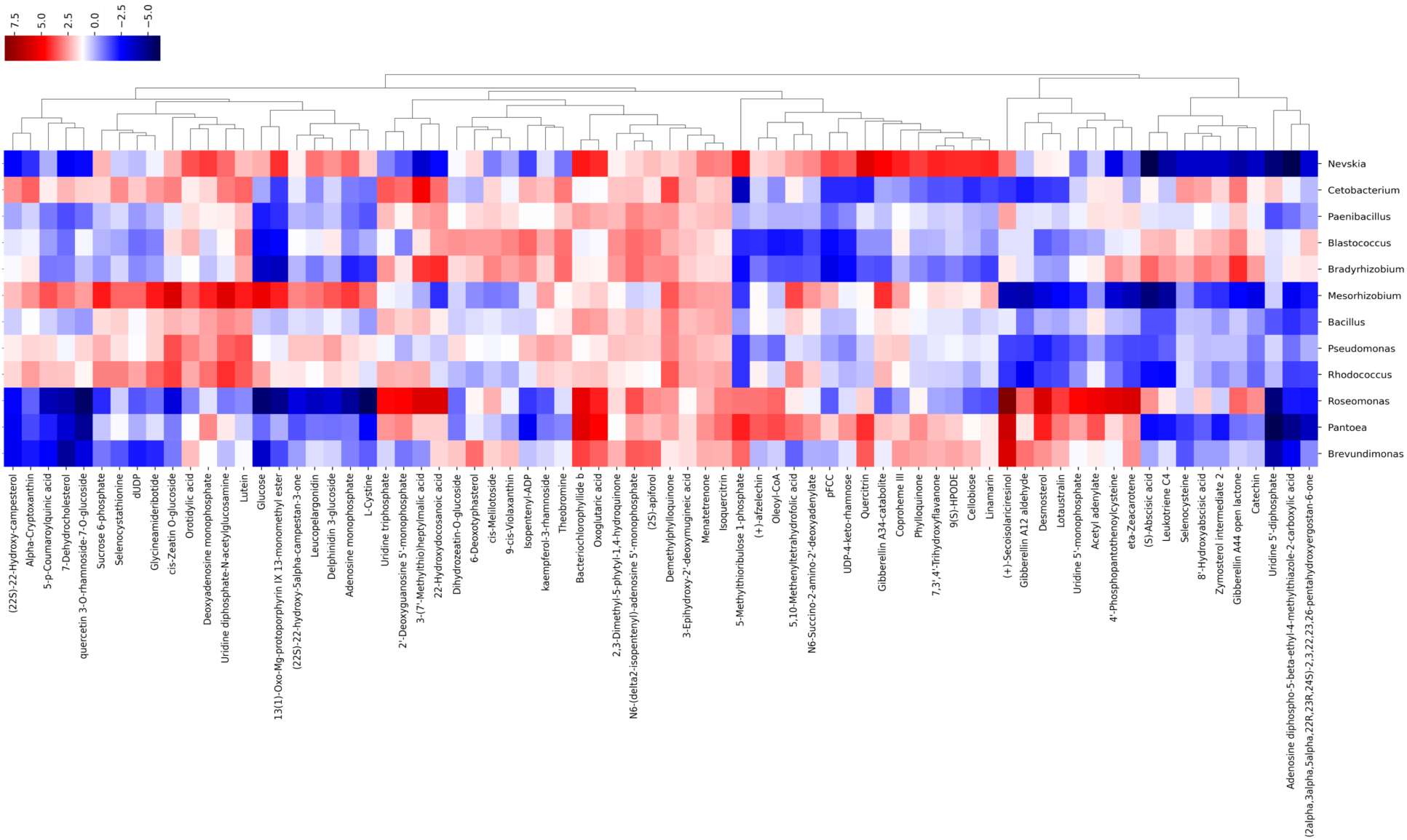
Associations between exudate compounds and bacterial genera inferred by the mmvec neural network.^33^ Colors indicate the estimated log conditional probability of observing a root exudate compound given the presence of a microbial genus. Root exudates are clustered based on their associations with microbial genera.

Here four flavonols, including three quercetin derivatives and one kaempferol derivative, had varying co-occurrence with the 12 bacterial genera: Isoquercitrin (quercetin 3-O-glucoside) had very high co-occurrence with all the 12 genera, particularly *Pantoea*; quercetin 3-O-rhamnoside-7-O-glucoside and kaempferol-3-O-rhamnoside had similar co-occurrence patterns with several microbes, opposite to quercitrin (quercetin 3-O-rhamnoside) (**Fig 5**). Quercetin and its derivatives can stimulate PGPR, including *Pantoea* that harbors quercetin dioxygenases for degrading flavonols.^45,71^ This aligns with our observation of very high co-occurrence probabilities between *Pantoea* and two quercetin derivatives (quercitrin > isoquercitrin). Three GAs, GA12-aldehyde (early precursor of bioactive GAs), GA44 open lactone (key intermediate in bioactive GA1 synthesis), and GA34-catabolite (deactivated catabolite from bioactive GA4), had varying co-occurrence with all the 12 bacterial genera, with GA44 open lactone and GA34-catabolite showing opposite patterns. GA phytohormones are crucial for many aspects of plant growth and development. Appropriate levels of bioactive GAs are tightly regulated in plants and also modulated by PGPR such as *Bacillus*, *Bradyrhizobium*, and *Mesorhizobium* that possess GA metabolic pathways.^72–74^ Here *Bacillus* co-occurred with GA34-catabolite, while *Bradyrhizobium* and *Mesorhizobium* had very high co-occurrence probability with GA44 open lactone and GA34-catabolite, respectively. (S)-Abscisic acid, the naturally occurring and bioactive form of phytohormone ABA, and its direct catabolite 8’-hydroxyabscisic acid had high co-occurrence with *Blastococcus*, *Bradyrhizobium*, and *Roseomonas*, suggesting their participation in ABA metabolism. Indeed, PGPR like *Bradyrhizobium* can regulate ABA levels in the rhizosphere through biosynthesis and catabolism of ABA.^41,72^ Two glycosides of phytohormone cytokinins, cis-zeatin-O-glucoside and dihydrozeatin-O-glucoside, had opposite associations with several PGPR, with cis-zeatin-O-glucoside showing very high co-occurrence with four PGPR (*Mesorhizobium* > *Rhodococcus* > *Pseudomonas* > *Bacillus*). Among these PGPR, *Rhodococcus* and *Pseudomonas* are known to produce cytokinins.^72,75^

Taken together, our analyses of hypothetical community growth rate, biofilm formation tendency, community assembly mechanism, and specific microbe-exudate associations suggest that biochar-induced root exudates may play a key role in the rhizosphere microbiome restructuring. In particular, secondary metabolites such as flavonoids (flavonols) and developmental phytohormones (GAs, ABA, cytokinins) may evoke diverse PGPR capable of utilizing these compounds. While biochar-induced microbiome shifts in bulk soil has been well documented (e.g., our meta-analysis^3^), this study suggests additional modulation mechanisms in the rhizosphere through metabolic coupling of root exudates with microbial substrate-use preferences. Future research employing metagenomics and metatranscriptomics, as well as using pure PGPR cultures, is required to ascertain our observations and elaborate underlying biochemical mechanisms.

### 3.4. Biochar-treated rhizosphere microbiome has shifted functions

#### 3.4.1. Shifted nitrogen and methane cycling

Many studies have reported biochar’s impact on N and methane cycling in bulk soil, partially due to changes in microbial communities.^2,3^ We hypothesized such effects also occur in the rhizosphere with additional influences from root exudates. Our qPCR measurements found increased abundance of *nifH* (N fixation), *nxrA* (nitrification, NO_2_^-^ to NO_3_^-^), *nirS* (denitrification, NO_2_^-^ to NO) and *nosZ* (denitrification, N_2_O to N_2_), and reduced abundance of *nirK* (denitrification, NO_2_^-^ to NO) in the biochar-treated rhizosphere, although the differences were insignificant (**Fig 6B**). These trends are consistent with the emergence of N-fixing bacteria (e.g., *Allorhizobium-Neorhizobium-Pararhizobium-Rhizobium*, *Azospirillaceae*) as keystone taxa and the enrichment of ammonia-oxidizing bacteria (e.g., *Nitrosomonas*) in the biochar-treated rhizosphere (**Fig 4C**; **Table S5**). Interestingly, biochar-induced increase in *nxrA* abundance coincided with a drastic enrichment of *Roseomonas*, a genus known to harbor *nxrA*,^76^ and showed high co-occurrence with biochar-induced ABA (a terpenoid phytohormone) and its direct catabolite 8’-hydroxyabscisic acid (**Fig 5**). In complete denitrification, NO_2_^-^ is reduced to NO by nitrite reductases (*nirK* and *nirS*), while NO is further reduced to N_2_O by nitric oxide reductases (*nor*) and N_2_O reduced to N_2_ by nitrous oxide reductase (*nosZ*). Changes in the relative abundance of these genes are associated with the net emission of N_2_O. Specifically, the (*nirK*+*nirS*)/*nosZ* ratio has been used to estimate N₂O emission potential.^77^ Our qPCR results showed a reduced (*nirK*+*nirS*)/*nosZ* ratio in the biochar-treated rhizosphere, suggesting a promotion of complete denitrification and a reduction of N₂O emission potential, likely due to biochar’s role in facilitating electron transfer to denitrifying microbes.^78^ qPCR results also showed a reduction in the (*nirK*/*nirS*) ratio, suggesting a shift from *nirK*-type nitrifiers, which often lack the *nosZ* gene, to *nirS*-type nitrifiers.^79^ This observation is consistent with the emergence of *Magnetospirillum* as keystone taxa and the enrichment of *Sporomusa* in the biochar-treated rhizosphere (**Fig 4C**; **Table S5**). *Magnetospirillum* harbors a complete denitrification pathway, able to reduce NO_3_^-^ to N_2_.^80^ It should be noted that certain *Sporomusa* species contain a recently-characterized novel lactonase-type N_2_O reductase that cannot be captured by existing primers including those used here.^81^ Overall, these results suggest that biochar may promote nitrogen fixation and nitrification while reducing N_2_O emission potential in the rhizosphere through restructuring the rhizosphere microbiome.

**Fig 6.**
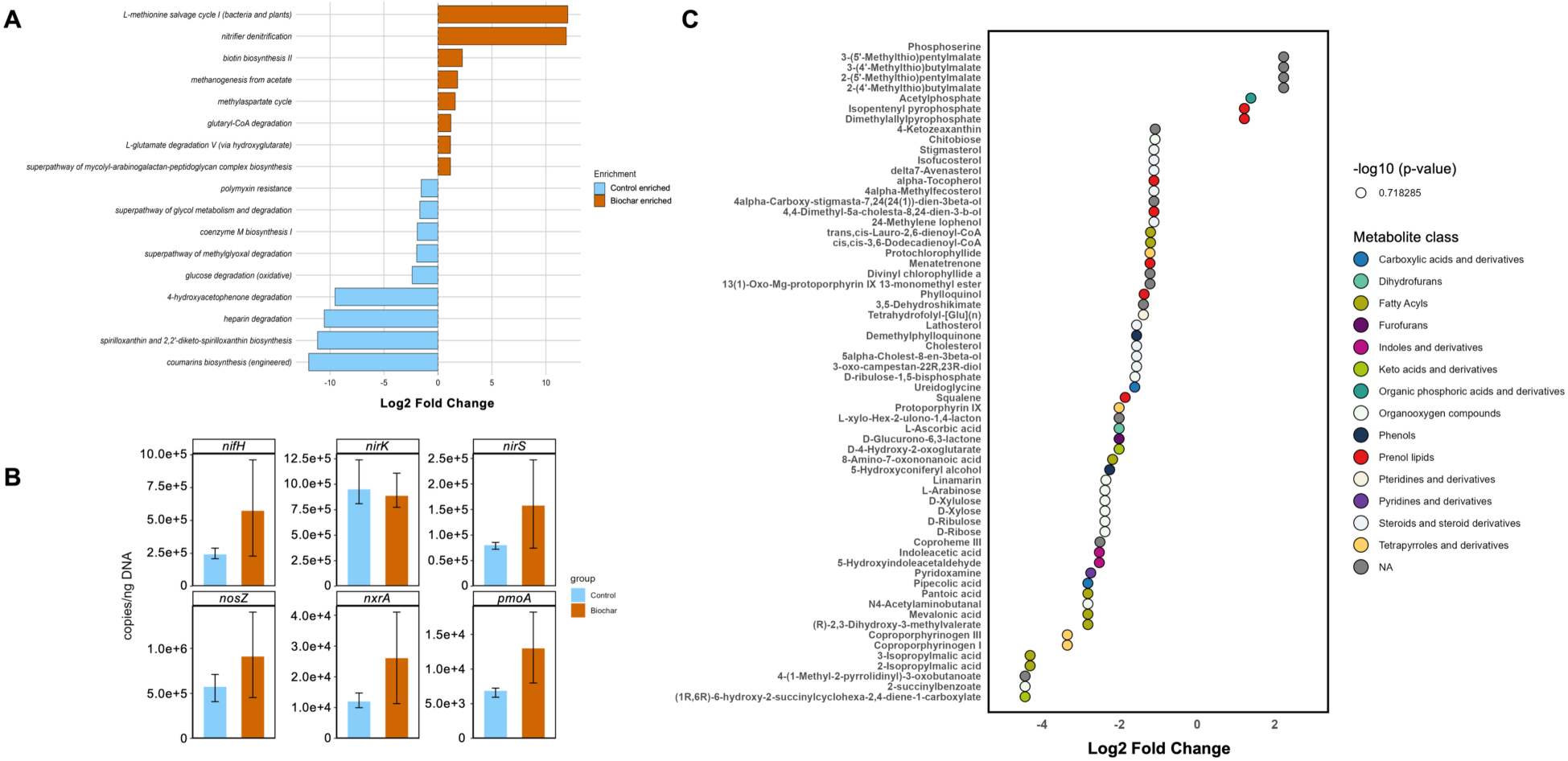
(A) Microbial pathways (from PICRUSt2^26^) significantly regulated by biochar (|Log_2_FC| > 1, *p-adjusted* < 0.05). (B) Abundance of key nitrogen- and methane-cycling genes from qPCR. Boxplots represent the median and interquartile range. (C) 64 most discriminative rhizosphere soil exometabolites influenced by biochar (|Log_2_FC| > 1, *p-adjusted* < 0.05). Compounds are grouped into ChemOnt classes.

qPCR did not detect *mcrA* encoding the methyl-coenzyme M (CoM) reductase complex in any sample, reflecting a low representation of *mcrA*-containing methanogens in the ectorhizosphere. Consistently, PICRUSt2 predicted that biochar suppressed CoM biosynthesis I, which is required to produce methyl-CoM (**Fig 6A**).^82^ qPCR found increased *pmoA* (encoding membrane-bound particulate methane monooxygenase^83^) in the biochar-treated rhizosphere, suggesting enhanced methane oxidation although the differences were insignificant (**Fig 6B**). Consistently, two C5 prenol lipids, isopentenyl pyrophosphate and dimethylallylpyrophosphate, were only detected in the biochar-treated soil metabolome (**Fig 6C; Table S12**). These prenol lipids are building blocks of all isoprenoids, including bacteriohopanepolyols that are crucial structural components of methanotroph membranes. Their detection in the biochar-treated soil metabolome may suggest stimulated growth and subsequent lysis of methanotrophs. Further, biochar may have influenced methylated substrates and thereby methylotrophic methanogenesis,^84^ as suggested by PICRUSt2-predicted enrichment (>1000 fold) of methionine salvage pathway (MSP) in the biochar-treated rhizosphere (**Fig 6A**) and the detection of several MPS intermediates in root exudates and the soil metabolome, as well as high co-occurrence of specific PGPR and a methylthio compound downstream in MSP (detailed in the SI). MSP produces methanethiol, a methylated substrate for methylotrophic methanogenesis; both methanethiol methylation and methionine also influence dimethylsulfide, another methylated substrate for methylotrophic methanogenesis.^84,85^ However, this hypothesis remains to be verified by metagenomic and metatranscriptomic analysis. The importance of methylotrophic methanogenesis in agricultural methane emissions has only recently been recognized.^86,87^ Given the large suite of methylated substrates in root exudates that may be utilized by methylotrophic methanogens and biochar’s impact on the chemistry of root exudates, more experimental and field research is needed in this direction.

Together, our results from qPCR, function prediction, and untargeted soil metabolomics suggest that in the rhizosphere, biochar-induced microbiome restructuring may lead to shifts in N and methane cycling, resulting in attenuated GHG emissions. Our results also highlight an urgent need of scrutinizing how biochar-induced changes in root exudate chemistry may affect methylotrophic methanogenesis.

#### 3.4.2. Broader changes in rhizosphere microbiome functions

A total of 506 exometabolites were detected in rhizosphere soils (**Fig 7A**). Biochar significantly influenced the soil metabolome chemistry (PERMANOVA: *p* = 0.04 for treatment) (**Fig 7B**), reflected by 64 most discriminative compounds from 14 ChemOnt classes (|Log_2_FC| > 1, *p-adjusted* < 0.05) (**Fig 6C**) (details in the SI). Remarkably, several plant defense compounds were only detected in the biochar-treated rhizosphere, including lotaustralin and methionine-derived methylthioalkylmalates that are important precursors of sulfur-containing glucosinolates (**Fig 6C; Table S12**).^41^ Further, IAA and an indole derivative (5-hydroxyindoleacetaldehyde) were detected in the control rhizosphere but absent in the biochar-treated rhizosphere where 5-hydroxyindoleacetic acid, a direct oxidation product of 5-hydroxyindoleacetaldehyde was detected (**Fig 6C; Table S3; Table S12**). This may suggest microbial metabolism of IAA and is consistent with the enrichment of IAA-producing PGPR (Section 3.3.1).

**Fig 7.**
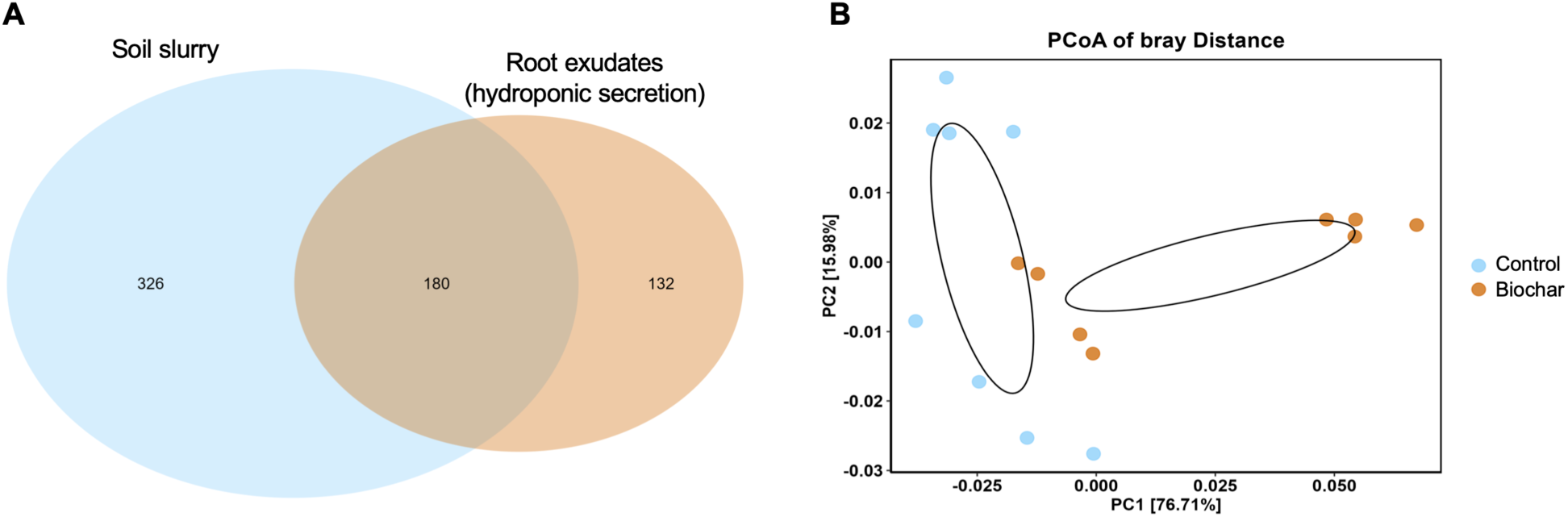
(A) Venn diagram showing root exudates from hydroponic secreted (312 compounds), metabolites in soils (506 compounds), and exometabolites including microbial products (326 compounds) there were detected under the biochar treatment. (B) PCoA of Bray-Curtis distance of soil metabolites detected in the control and biochar-treated rhizosphere (PERMANOVA test: *p* = 0.039 for experiment round; *p* = 0.040 for treatment). Data represents the results from two rounds of independent experiments. The ellipses represent 95% confidence.

Among the 506 exometabolites, 180 compounds were detected in root exudates collected by hydroponic secretion, while 326 compounds were only found in the soil metabolome (**Fig 7A; Table S12**). The 326 exometabolites consisted of compounds secreted by roots in contact with rhizosphere microbes, as well as microbial metabolites. DIBOA-Glc was only detected in hydroponic secretion (**Fig 2B**), while its active form DIBOA was only detected in soil exometabolites and had high co-occurrence probability with 7 PGPR (**Fig S14**), suggesting BXs may be used as signaling cues for recruiting PGPR.^40,75^ Various GAs were detected in both root exudates and soil exometabolites, but an early precursor of bioactive GAs (GA12-aldehyde) was only detected in root exudates, whereas two GA catabolites (GA8, GA8-catabolite) and one intermediate (GA24) were only detected in the soil metabolome (**Table S12**), with GA24 showing high co-occurrence probability with several PGPR (**Fig S14**), especially *Bradyrhizobium* that is known to possess GA metabolic pathways.^72–74^

After mapping the 326 soil exometabolites to predicted microbial pathways (**Table S13**) and comparing microbe-root exudate and microbe-exometabolite co-occurrence networks (**Table S9**), we found biochar likely influenced microbial synthesis of three vitamins (biotin, phylloquinone, menaquinol) (more details in the SI). Menaquinone is a vitamin (K12) produced by bacteria and some archaea. Here, the menaquinol percussor 2-succinyl-5-enolpyruvyl-6-hydroxy-3-cyclohexene-1-carboxylic acid was only detected in root exudates, while two immediate metabolites of menaquinone biosynthesis, (1R,6R)-6-hydroxy-2-succinylcyclohexa-2,4-diene-1-carboxylate and 2-succinylbenzoate, were only detected in the soil metabolome (**Table S3; Table S12**). Biochar significantly reduced the two intermediates (**Fig 6C**). Coincidentally, PICRUSt2 predicted that biochar increased all but one menaquinone biosynthesis genes; meanwhile, menaquinol, the bioactive reduced form of menaquinone, was detected in the soil metabolome (**Fig S15**). These results may suggest biochar-induced boost in microbes that use cellular menaquinone pool for respiratory electron transfer.^88,89^ Whether or not these microbes contribute to more complete denitrification, as well as other redox reactions, remains to be investigated.

Collectively, our results from untargeted soil metabolomics and function prediction suggest profound modulating effects of biochar on rhizosphere microbiome functionality, ranging from phytohormone metabolism by PGPR to biosynthesis of redox-active quinone compounds. These changes could have significant implications for plant growth and soil biogeochemistry. Future research should integrate metagenomic, metatranscriptomic, and metabolomic data to validate our observations and elaborate underlying microbial mechanisms.

### 3.5. Implications for sustainable crop production and future needs

Rhizosphere engineering has attracted increasing interest as a promising approach for sustainable crop production.^90^ Extending from our prior work on metabolites within root tissues,^5^ this study is among the first that elaborates biochar-induced systematic shifts in secreted root exudates, the rhizosphere microbiome, and the soil metabolome. Our results suggest new chemical and microbial mechanisms behind the widely reported but poorly understood effects of biochar on the soil-plant system. Specifically, biochar evokes a plant-beneficial rhizosphere microbiome centered by diverse PGPR, and such microbiome restructuring may partially be attributed to biochar-induced changes in root exudate chemistry, particularly complex molecules including a wide range of plant secondary metabolites. Therefore, compared to its modulating effect on the bulk soil microbiome, biochar also modulates the rhizosphere microbiome indirectly through altering rhizosphere chemistry. The restructured rhizosphere microbiome shows alterations in N and methane cycling that may lead to reduced potential of N_2_O and CH_4_ emissions. Additionally, the restructured microbiome is likely involved in the transformation of signaling cues and a variety of redox reactions, thus having broader implications for the soil-plant system.

Several limitations of this study should also be noted. First, the EcoFAB device and the cultivation condition simplify the complexity of the soil-plant system in root exudate chemistry, rhizosphere microbiome diversity, and root-microbe interactions. Future research should leverage larger-scale laboratory experiments and field studies and use other plant models to validate and extend findings from this study. Second, we identified potential associations between specific root exudate compounds and PGPR using a combination of network analysis and machine learning. These associations and the underlying biochemical mechanism should be ascertained by integrated metagenomic/metatranscriptomic analysis and in experiments using model bacterial strains. Third, we used 16S rRNA gene sequencing-based function prediction coupled with untargeted soil metabolomics to infer microbiome functions. Integrating metagenomic, metatranscriptomic, and metabolomic data will provide more conclusive evidence for biochar-induced community-level functional shifts and the links to biochar-induced changes in root exudate chemistry. Nevertheless, this research provides new insights into biochar’s multifaceted roles in reprogramming the rhizosphere and highlights the potential of engineering the rhizosphere through reshaping root-microbe interactions using designed biochar or root exudate cocktails. Further, the use of crop residue-derived biochar also exemplifies a circular economy approach valorizing agricultural wastes.^91^

---

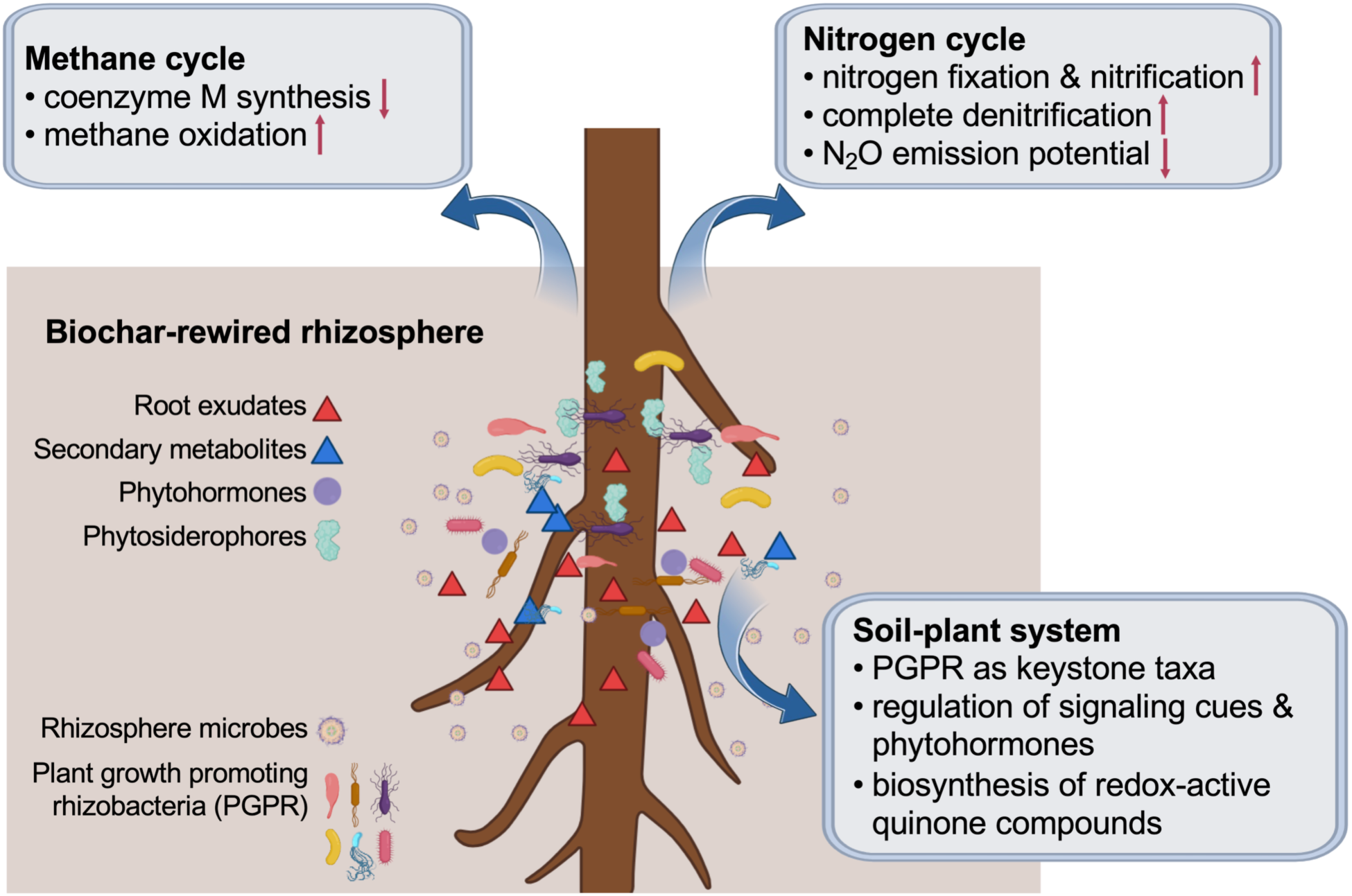
**Graphic TOC**

## DATA AVAILABILITY STATEMENT

Raw reads are available at the NCBI Sequence Read Archive under the BioProject accession number PRJNA1394313. Supporting Information tables (Table S1-S13) are also available at https://doi.org/10.5281/zenodo.18930558.

## SUPPORTING INFORMATION

Additional methods (biochar production and characterization, EcoFAB fabrication, plant experiment and sample collection, untargeted metabolomics, next generation sequencing and data processing, qPCR and data normalization, biofilm assay, multi-omics integration, general statistics and data visualization), and results and discussion (characterization of wheat biochar, biochar-induced differential root exudation, rhizosphere microbiome restructuring and its association with biochar-induced root exudates, biochar-treated rhizosphere microbiome with shifted functions) (PDF).

Properties of the agricultural soil and wheat biochar, qPCR primers and conditions, compounds detected by Orbitrap HRMS, compounds detected in root exudates but not inside of root tissues, microbes enriched in the biochar-treated or control rhizosphere, topological properties of microbial co-occurrence networks reconstructed with SparCC, keystone genera identified by centrality metrics (degree, betweenness, and closeness), keystone genera idefined by within-module connectivity and between-module connectivity, predicted microbial pathways significantly associated with biochar-induced root exudates, PICRUSt2 predicted N-cycling gene abundance, PICRUSt2-predicted top 5 genera carrying denitrification genes (*narG*, *nirK*, *nirS*, *nosZ*) and their relative abundance, metabolites detected only in rhizosphere soils and not in root exudates, predicted microbial pathways significantly associated with biochar-induced soil exometabolites (XLSX).

## AUTHOR INFORMATION

### Author Contributions

**Hanyue Yang:** Investigation, Methodology, Data curation, Formal analysis, Validation, Visualization, Writing – original draft, Writing – review and editing. **Amina F. Mughal:** Methodology, Formal analysis. **Yaqi You:** Conceptualization, Funding acquisition, Resources, Project administration, Supervision, Formal analysis, Validation, Visualization, Writing – review and editing.

### Funding

This work was supported by the State University of New York College of Environmental Science and Forestry (SUNY ESF) through startup funds to Yaqi You. It was partially supported by the US Department of Agriculture through a McIntire-Stennis Capacity Grant (NI23MSCFRXXXG070) and the State University of New York Empire Innovation Program through its Chancellor’s Early Career Scholar program. Hanyue Yang is grateful to the ESF Alumni Association.

### Notes

The authors declare no competing financial interest.

## ACKNOWLEDGEMENTS

We thank the group of Dr. Amin Mirkouei (University of Idaho) for producing the biochar used in this study. We would like to posthumously acknowledge Kevin Guerin (ESF Analytical and Technical Services) for great assistance with fabricating acrylic mold parts on a lathe machine. We thank Dr. Alex Artyukhin for training H. Yang on the Q Exactive HF Orbitrap mass spectrometer. We thank undergraduate students Olivia Abry and Hayden Bregg (Department of Biology, ESF) for assistance with the EcoFAB experiment, and Dr. Leanne Powers (Department of Chemistry, ESF) for assistance with DOC measurement. We are grateful to Syracuse University Research Computing for providing high-performance computing resources.

## Notes

https://doi.org/10.5281/zenodo.18930558

https://www.ncbi.nlm.nih.gov/bioproject/PRJNA1394313/

### Competing Interest Statement

The authors have declared no competing interest.

### Summary of Updates

Title and text (Abstract, Results and Discussion) updated to improve data interpretation; figures updated to improve data presentation; Supplemental file (PDF) updated.

