## Supplementary Information for "Biochar Rewires Root Exudates and the Rhizosphere Microbiome and Its Functionality"

*Correspondence:

Yaqi You

Number of pages: 36

Number of figures: 15

Number of tables: 13 (separate XLSX file, also available at doi:10.5281/zenodo.18930558)

### 1. SUPPLEMENTARY MATERIALS AND METHODS

#### 1.1. Biochar production and characterization

Biochar was generated by slow pyrolysis of wheat straw at 350 °C for 30 minutes under a no-oxygen condition, the same as in our prior study.[1,2] Biochar was characterized using conventional and advanced approaches.[1] Regular characterizations, including pH, major and trace elements, nitrogen (N) species (i.e., $NH_4^+$-N and $NO_3^-$-N), organic matter (OM), electrical conductivity (EC), and cation exchange capacity (CEC) were performed at the Environmental Analytical Laboratory of Brigham Young University (Provo, Utah, USA) using conventional soil methods like before (**Table S1**). Advanced characterizations, including scanning electron microscopy (SEM) for microscopic structure, energy-dispersive X-ray spectroscopy (EDS) for surface elemental composition, Fourier-transform infrared spectroscopy (FTIR) for molecular composition and functional groups, and X-ray diffraction (XRD) for crystal structure, were performed at the Microscopy and Characterization Suite, Center for Advanced Energy Studies (CAES) and in the Department of Chemistry, Idaho State University. FTIR and XRD data analysis was performed in R using the R package "MALDIquant" and "baselineWavelet".[3,4] Boehm titration was performed at SUNY ESF to measure three oxygen-containing functional groups (acidic carboxyl, lactone, and phenolic group) on the surface.[1]

#### 1.2. EcoFAB fabrication

Design files were downloaded from the official website (https://eco-fab.org/device-design/),[5] modified using the autoCAD software, and uploaded to a MakerBot Replicator 2X 3D printer to fabricate the mold frame (172.8 (L) × 140.80 (W) × 25.00 (H) mm) with acrylonitrile butadiene styrene (ABS) filament (**Fig 1A**). Acrylic mold parts, including a rectangle base (172.0 (L) × 140.0 (W) × 6.0 (H) mm) and an oval chamber (61 (L) × 40 (W) × 2.5 (H) mm) were fabricated on a lathe machine (**Fig 1A**). We chose acrylic over ABS for these mold parts due to the need of smooth interior wall surfaces for the EcoFAB chamber. These parts were assembled into a complete mold with an oval chamber (97.6 (L) × 64 (W) × 3.2 (H) mm) for the casting of a polydimethylsiloxane (PDMS) layer (**Fig 1A**).

The SYLGARD 184 silicone elastomer kit (Dow Inc.) was used to prepare PDMS materials. After optimization, we chose a base to curing agent ratio of 25:1 to achieve the most appropriate PDMS stiffness. Silicon elastomer and curing agent were thoroughly mixed, degassed in a vacuum chamber for at least 30 minutes, and poured into the complete mold (**Fig 1A**). Air bubbles were allowed to rise to the surface and removed by blowing the PDMS surface with compressed air, after which the mold was heated to 85 °C in a heating block and the crosslinking density was continuously monitored. Afterwards, extra edges of the PDMS layer were carefully removed.

The surfaces of the PDMS layer and a 178 (L) × 127 (W) mm microscope slide (Ted Pella, Inc. #260234-25) were rinsed with pure methanol, blow-dried with compressed air, and quickly and firmly adhered to each other to form the EcoFAB device (**Fig 1A**). To maintain a sterile environment for plant experiments, each EcoFAB device was enclosed in a transparent cover box, and the whole system was sterilized using 70% ethanol for 30 minutes and then 100% ethanol for 5 min before being used in the plant experiment (**Fig 1B**).

##### 1.3. Plant experiment and sample collection

Wheat (*Triticum aestivum* L.) seeds were soaked in 70% ethanol for 2 minutes, rinsed with sterile Milli-Q water for three times, soaked in 10% bleach (sodium hypochlorite) for 5 minutes, and rinsed with sterile Milli-Q water for three times. Afterward, the seeds were incubated for 3 days at 25 °C and in the dark in a petri dish containing sterile Milli-Q water. After the seeds were germinated, seedlings of similar sizes were chosen and transferred aseptically into sterilized EcoFAB devices (one plant per device) pre-filled with soil slurries with or without biochar and cultivated for 21 days under controlled light (12 hour of full spectrum LED light, 12 hour in dark) and humidity (80% by refilling Milli-Q water every three days) conditions.

After 21 days, individual EcoFAB devices were dissembled, and the remaining soil slurries and plant seedlings were collected in sequential order: (1) rhizosphere soil slurry for the soil metabolome, (2) ectorhizosphere soil for DNA extraction and DNA-based analyses, and (3) after removing ectorhizosphere soils, 24 hour hydroponic secretion to collect root exudates (**Fig 1C**).

Soil metabolite extraction followed procedures similar to the extraction of root exudates (detailed in the main text). An aliquot of 1.5 mL of a soil slurry sample was added to a 2.0 mL microcentrifuge tube, dried using a vacuum concentrator (Savant SpeedVac SPD300DDA, Thermo Scientific), and 1.5 mL of methanol in Milli-Q water (80% v/v) was added to the tube, followed by vortexing for 15 min and sonication for 5 min at room temperature. The resulting extract was filtered through a 0.22 μm PVDF filter (Millipore) and stored at -80 °C until untargeted metabolomics analysis.

##### 1.4. Untargeted metabolomics

Root exudates and soil slurries were analyzed using liquid chromatography-mass spectrometry (LC-MS).[6] For LC, the mobile phase consisted of 0.1% (v/v) formic acid in Milli-Q water (solvent A) and 0.1% formic acid in LC/MS grade acetonitrile (solvent B). The gradient elution program was set as follows: 5% B (0–1.5 min), linearly increased to 100% B (1.5–12.5 min), maintained at 100% B (12.5–18.0 min), then decreased to 5% B (18.0–18.5 min), and finally maintained at 5% B for column re-equilibration until the end of the run at 20.0 min. The flow rate was maintained at 300 μL/min. The MS conditions were as follows: electrospray ionization (ESI) in positive ion (ESI+) and negative ion mode (ESI-) with a spray voltage of 3700V and -4000V, respectively, endplate offset of -500V, nebulizer gas pressure of 15 psi, dry gas flow rate of 4.0 L/min, dry gas temperature of 180 °C, and mass range of 20-1200 m/z. For untargeted metabolomics, a full scan of data acquisition was applied. The full MS scan was performed with a resolution of 240,000 (at 200 m/z) across a mass range of 20-1200 m/z. The automatic gain control (AGC) target was set to 1E6 with a maximum injection time of 500 ms.

Principal coordinates analysis (PCoA) was performed based on Jaccard distance using the "vegan" package.[7] For network analysis, pairwise Pearson correlation with false discovery rate (FDR) control was calculated using the "WGCNA" package;[8] only significant correlations ($r$ > 0.9 and adjusted $p$ < 0.01) were retained. Network topology was assessed with the "igraph" package.[9] Node degree distributions were fitted to a power-law model, and nodes with degrees exceeding the 95$^{th}$ percentile were identified as highly connected hub nodes.[1]

##### 1.5. Next generation sequencing and data processing

For 16S rRNA gene sequencing, the V4 region of the prokaryotic 16S rRNA gene was amplified with the 515F/806R primer pair.[10,11] After cleanup, size assessment, and quantification, libraries were pooled in equimolar concentrations and sequenced on an Illumina MiSeq platform (2×250 bp) at Idaho State University Molecular Research Core Facility.

Raw reads were processed using QIIME2, DADA2, and “phyloseq”: Paired-end reads were subjected to Quality Assurance/Quality Control, trimmed at both ends when the Phred score fell below 33, and denoised with DADA2 to yield amplicon sequence variants (ASVs).[1,12–14] Rare ASVs (less than 5 occurrences across all samples or present in only one sample) were filtered out to avoid sequence errors and artifacts; taxonomy was assigned using a naive Bayes machine-learning classifier trained for the 515F-806R V4 region against the SILVA database (release 138); unassigned reads and reads associated with Eukarya, mitochondria, and chloroplast were removed.[15] Alpha diversity was assessed at the ASV level after rarefaction. Beta diversity was estimated based on the Bray-Curtis distance after the log-transformation of relative abundance or the Weighted UniFrac distance using the ASV abundance.

When predicting microbial functions using PICRUSt2 analysis, any ASV having a nearest-sequenced taxon index (NSTI) greater than two was filtered out.[16] Enzyme Commission (EC) numbers and KEGG orthologs (KOs) were used to estimate the abundance of gene families. The abundance of metabolic pathways was estimated by mapping EC gene families based on the MetaCyc database,[17] and significant changes were identified using methods (e.g., DESeq2) implemented in the “ggpicrust2” R package ($p < 0.1$).[18,19] For a sample, the relative abundance of a KO was calculated using the following equation, which can be directly compared to the normalized qPCR result: Relative abundance of the $i$th KO = total reads of the $i$th KO / total reads of all ASVs.

For network analysis, we used SparCC with a $p$-value determined by bootstrapping (1000 times).[20] Significant correlations ($p < 0.05$) were retained. We identified keystone nodes based on two criteria: (1) Nodes that were above the 95% quantile of the fitted log-normal distribution of three centrality metrics (degree, betweenness, and closeness);[21] or (2) nodes that were network connectors or module hubs.[22,23]

**1.6. qPCR and data normalization**

While PICRUSt2 inference provides a more comprehensive view of community functions, the inference accuracy varies depending on samples. qPCR was used to quantify the abundance of bacterial 16S rRNA gene and genes related to N and methane cycling using established primers and PCR conditions listed in **Table S2**. The thermal program consisted of an initial denaturation at 95°C for 2 min, followed by 35 (16S, *mcrA*, *nxrA*) or 40 (*nifH*, *nirK, nirS*, *nosZ*, *pmoA*) cycles of 10 sec at 95 °C, 20 sec at the annealing temperature (**Table S2**), and 30 sec at 72 °C. A melting curve analysis was conducted after each run to verify reaction specificity. Standard curves were generated using a cloned PCR product for 16S rRNA gene and synthetic DNA (gBlocks, Integrated DNA Technologies) for other genes (**Fig S1**), following the MIQE guidelines.[24] Negative control was included in each run. A melt curve analysis was conducted after each qPCR run to verify reaction specificity. The PCR product was denatured (95°C for 2 min) and annealed (at temperature in **Table S2** for 1 min) before melting curve analysis, which consisted of increasing the temperature at 0.5°C/10 sec to 95°C. Standard curves (2 to 9 log10 range in duplicate) were generated using cloned PCR product for 16S rRNA gene and synthetic DNA (gBlocks, Integrated DNA Technologies) for other genes:[25,26] (1) 2233 bp single strand DNA containing 5 N-cycling genes and an internal reference (142 bp fragment of Zika virus that is rare in soils), and (2) 1213 bp single strand DNA containing 2 methane cycling genes and the same internal reference (**Fig S1**). We chose these gene fragments because they reflect amplicons from reference organisms using primers listed in **Table S2**, and because their feasibility has been proven for terrestrial ecosystems.[25] Correlation coefficients from standard curves reached $R^2 > 0.96$ for *nifH*, *nirK* and *pmoA*, and $R^2 \geq 0.99$ for others, with amplification efficiencies of 84% to 110%. Detection limits were <14, <25, <35, <304 copies for bacterial 16S rRNA gene, N-cycling genes (*nifH*, *nxrA*, *nirK*, *nirS*, *nosZ*), *mcrA*, and *pmoA*, respectively. Gene abundance was normalized against DNA concentration to calculate copies of each gene per ng DNA. We chose not to normalize the abundance of functional genes against total copies of 16S rRNA gene because the 16S rRNA gene primers used here were designed only for eubacteria,[27] while microorganisms other than bacteria can also harbor these functional genes.

**1.7. Biofilm assay**

*Pseudomonas putida* KT2440, a soil bacterium, was used to test whether biochar-induced root exudates can promote biofilm formation compared to root exudates from control plants.[28] The strain was grown on Luria-Bertani (LB) agar or in LB broth at 30°C overnight. Biofilm assay was conducted in 96-well microtiter plates (Corning) as others.[29] Each well contained 300 μL root exudates from either biochar treatment or control and 3 μL of the overnight *P. putida* KT2440 culture or no-bacteria control. Three technical replicates were included for each root exudate sample and no-bacteria control, respectively. Plates were incubated at 27°C for 48 hours. Then wells were washed with phosphate-buffered saline (PBS) solution (1 mM $KH_2PO_4$, 1 mM $Na_2HPO_4$, 2.7 mM KCl, 13.7 mM NaCl, pH 7.4) for three times to remove any unbound cells, and biofilms were stained with 300 μL of 0.1% crystal violet solution for 20 minutes and washed with PBS to remove excessive crystal violet. After adding 300 μL of acetone-ethanol mixture (20:80, v/v) to each well, optical density at 540 nm ($OD_{540}$) was measured on a BioTek Synergy H1 hybrid microplate reader (Agilent Technologies) to quantify biofilm mass. $OD_{540}$ readings from the blank wells were averaged and subtracted from the readings of the wells containing biofilms.

Dissolved organic carbon (DOC) content of root exudates, measured as non-purgeable organic carbon (NPOC), was determined using a Shimadzu total organic carbon analyzer (TOC-VCPH) with high-temperature catalytic combustion (720°C) and non-dispersive infrared (NDIR) detection. Collected root exudates were filtered through a 0.22 μm PVDF membrane filter. Filtrate was diluted by 10-fold with Milli-Q water, acidified with 2N HCl, and purged with an inert gas to remove inorganic carbon and volatile organic compounds. The remaining non-volatile organic carbon was quantified on the TOC analyzer.

##### 1.8. Multi-omics integration

To identify co-occurring root exudates and rhizosphere microbes, we used two approaches, OmicsNet 2.0 and mmvec. OmicsNet 2.0 utilizes the AGORA collection of high-quality genome-scale metabolic models to predict the metabolic capacity of microbial genera and pertinent proteins.[30,31] Potential score was set to 0.8, and metabolites lacking pathway annotation, currency metabolites, and universal metabolites were excluded. The KEGG database was used to build the network of root exudates and microbial proteins,[31] as we hypothesized that co-occurrence of root exudates and microbial genera were due to the processing of root metabolites by those genera. The neural network-based mmvec (microbe-metabolite vectors; https://github.com/biocore/mmvec) was also used.[32] Compared to approaches based on correlation (e.g., Spearman's) or network (e.g., SparCC), mmvec estimates the conditional probability of observing a metabolite given that a microbe is present. It can identify previously undiscovered patterns and is less prone to false discovery (positive or negative) as it provides consistency between absolute and relative abundances of compositional data. However, it does not directly access the statistical significance of an interaction inferred using co-occurrence probabilities, nor does it calculate confidence intervals for the strength of each metabolite-microbe interaction. We adapted the original Python script developed by Morton et al. (2019)[32] and implemented a one-hidden-layer neural network. Paired metabolomics and 16S rRNA sequencing datasets from the 16 samples were split into a training set (75%) and a testing set (25%), and model training parameters were set as default. Specifically, the model uses Songbird (https://github.com/biocore/songbird) to calculate the Aitchison distance between metabolites or between microbes,[33] and ordinates metabolites in microbe space. The heatmap visualization was implemented using matplotlib as in Morton et al. (2019).[32] Only those bacterial genera identified by both OmicsNet 2.0 and mmvec are shown in the heatmaps to ensure true positivity.

##### 1.9. General statistics and data visualization

Statistical analysis was conducted in R (v 4.2.1) and Python (3.7). For wheat biomass, significant changes between treatment and control were assessed using the Welch's *t*-test; for microbial alpha diversity, the nonparametric Wilcoxon test or Kruskal-Wallis test was used; for microbial beta diversity, permutational multivariate analysis of variance (PERMANOVA) was used.[34] Data visualization was done using the R packages "pheatmap", "VennDiagram", and "ggplot2", as well as the Python module "matplotlib.pyplot".[35–38] Cytoscape (v 3.9.1) and Gephi (0.10.1) were used for network visualization.[39,40]

For qPCR data from biological and technical replicates, to minimize the influence of outliers, only values within the interquartile range (IQR; 25th to 75th percentile) were retained for each gene. Statistical differences between treatment and control were tested using the permutation-based Wilcoxon test. Adjusted *p*-values were calculated using the Benjamini-Hochberg correction method. Boxplots were generated to visualize the median and IQR of gene abundances between groups. Similar approaches were also used for relative abundance of KOs predicted by PICRUSt2.

#### 2. SUPPLEMENTARY RESULTS AND DISCUSSION

##### 2.1. Characterization of wheat biochar

FTIR was used to characterize biochar molecular composition and functional groups. FTIR spectra showed an absorbance peak at 3008.34 $cm^{-1}$ attributed to the stretching vibration of =CH bonds in aromatic rings or olefins, and a peak at 2364.74-2322.46 $cm^{-1}$ attributed to –C≡N (**Fig S2**). Peaks ranging from 2000 to 400 $cm^{-1}$ were dominantly attributed to bending vibration (**Fig 2B**). Carboxyl C=O or aromatic C=C stretching vibration likely caused the 1742.59 $cm^{-1}$ peak, while –CH– bending vibration caused the 1371.11 $cm^{-1}$ peak. These peaks are generally considered characteristic of conjugated ketone and quinone groups. The peaks at 1277.49 $cm^{-1}$,

1262.39 $cm^{-1}$, and 1217.08 $cm^{-1}$ corresponded to C–O stretching vibration in esters or ethers, C–H bending vibrations in aromatic compounds, and possibly C–N or P–O vibration, respectively. Peaks at approximately 1000-500 $cm^{-1}$ are generally attributed to silicate/carbonate (**Fig 2B**).[41]

XRD was used to characterize crystal structures. XRD spectrum showed the presence of amorphous C ($2\theta = 24.6°$), graphite ($2\theta = 26.8°$), fluorite ($2\theta = 28.5°$), pyrophyllite ($2\theta = 29.5°$), and barite ($2\theta = 40.7°$) in biochar (**Fig 2D**).[42]

##### 2.2. Biochar induces differential root exudation

As shown in **Fig S4**, PCoA of Jaccard distance of 312 root exudates detected across samples from two independent rounds of EcoFAB experiments not only showed the influence of treatment on root exudation (PERMANOVA: $p = 0.145$ for round x treatment), but also demonstrated the heterogeneity of the biochar material, which could have caused variations between experimental rounds.

In our prior pot study, we detected 953 metabolites in root tissues of wheat plants grown under 0.25% wheat biochar.[1] In this EcoFAB study, 254 of the 953 root endometabolites (26.7%) were detected in hydroponically collected root exudates. This was expected as only a portion of root endometabolites are secreted. Two ChemOnt classes (benzene and substituted derivatives, pyridines and derivatives) were only detected in biochar-treated root endometabolites but not in root exudates (**Fig S5**). Compounds in these two classes are involved in aromatic compound metabolism (e.g., phenylalanine metabolism, tyrosine metabolism) and pyrimidine metabolism (e.g., nicotinate and nicotinamide metabolism, pyridine nucleotide biosynthesis), which are related to biosynthesis of numerous secondary metabolites. The non-detection of these two classes in biochar-treated root exudates may reflect their transformation to end-point products through the aforementioned metabolic pathways in the root and subsequent secretion.

It should be noted that while we carefully washed root tissues and used hydroponic secretion to collect root exudates in order to avoid microbial metabolites as much as possible, several metabolites detected in root exudates, including beta-1,4-poly-N-acetyl-D-glucosamine (chitin), bacteriochlorophyll *a*, geranylgeranyl bacteriochlorophyllide *a*, and geranylgeranyl bacteriochlorophyllide *b*, were from microbial sources (**Fig 3B; Table S3**). These compounds may be produced by endosphere microbes (i.e., microbes living inside roots) or taken up by roots from the rhizosphere before hydroponic secretion.

#### 2.3. Rhizosphere microbiome restructuring and its association with biochar-induced root exudates

##### *2.3.1. Biochar rewires the rhizosphere microbiome and microbial interactions*

In the native wheat rhizosphere, Proteobacteria, Cyanobacteria, Actinobacteria, Bacteroidota, Firmicutes, Planctomycetota, Verrucomicrobiota, Acidobacteriota, Myxococcota, and Bdellovibrionoya were the top 10 phyla (in descending order), comprising more than 60% of the microbiome and reflecting the core microbiome of plant rhizosphere (**Fig 4B**).[43] The top 40 genera in the wheat rhizosphere accounted for 35% of all prokaryotes (**Fig S7A**), and most of these genera are part of the core rhizosphere microbiome of cereal plants.[21]

Twelve genera were enriched by biochar: *Nitrosomonas* (Proteobacteria phylum), *Stella* (Proteobacteria phylum), and those in the Acidobacteriota phylum (*Subgroup_22*, *Elev-16S-1166*), the Cyanobacteria phylum (*SepB-3*, *Pantalinema*), and the Firmicutes phylum (*Fictibacillus*, *Fonticella*, *Oxalophagus*, *Solibacillus*, *Sporomusa, Sporacetigenium*) (**Fig S7B**; **Table S5**). Biochar also suppressed nine genera: *Actinoplanes* (Actinobacteriota phylum), *Candidatus_Yonathbacteria* (Patescibacteria), *Tolypothrix* (Cyanobacteria), and those in the phylum Proteobacteria (*Variovorax*, *Verticiella*), and the phylum Bacteroidota (*Nibribacter*, *Chryseobacterium*, *Emticicia*, *Rurimicrobium*) (**Fig S7B**).

As shown by microbial co-occurrence networks, biochar enhanced microbial interactions in the rhizosphere (**Fig 4C**; **Fig S8; Table S6-S8**). Keystone genera were identified based on two criteria: (1) Nodes that were above the 95% quantile of the fitted log-normal distribution of three centrality metrics (degree, betweenness, and closeness); (2) nodes that were network connectors or module hubs. Using the first criterion, we identified 36 and 43 keystone genera in the native and biochar-treated rhizosphere, respectively, including 10 shared keystone genera (**Fig S8**; **Table S7**). Using the second criterion, we identified 28 and 17 keystone genera in the native and biochar-treated rhizosphere, respectively, including 2 shared keystone genera (**Fig S8**; **Table S8**). Together, the native and biochar-treated rhizosphere microbiome shared twelve keystone genera, including *Blfdi19*, *Bradyrhizobium*, *Cellvibrio*, *Lysobacter*, *Microvirga*, *NS9_marine_group*, *Paracoccus*, *Rhodococcus*, *Shinella*, *Stenotrophomonas*, *Candidatus_Captivus*, and *Solirubrobacter*. Nine of them are plant-growth promoting rhizobacteria (PGPR) as described in the main text. The other three shared keystone genera have typical rhizosphere traits. For instance, *NS9_marine_group* bacteria (Flavobacteria phylum) are highly specialized in breaking down complex organic compounds like root exudates and can support other heterotrophic microbes through cross-feeding, thereby reinforcing microbial network complexity.[44] Besides the twelve shared keystone genera, the native rhizosphere also had other keystone genera with fitness advantage in the rhizosphere environment: *WD2101_soil_group* (Planctomycetes phylum) is able to degrade polysaccharides derived from plant cell wall or bacteria; acetogen *Sporomusa* (Firmicutes phylum) produces acetic acid that can be utilized by methanogens;[45,46] PGPR *Paenibacillaceae* (Firmicutes phylum) can improve plant growth by producing siderophore and phytohormone indole-3-acetic acid (IAA) (**Fig 4C**).[47]

###### ***2.3.2. Rhizosphere microbiome rewiring is associated with biochar-induced root exudates***

Multiple lines of observations from hypothetic community growth rate, biofilm formation tendency, and community assembly mechanism suggest that biochar-induced root exudates may play a key role in the rhizosphere microbiome restructuring, likely through metabolic coupling of root exudates with microbial substrate-use preferences. For example, biochar upregulated exudation of 2,4-dihydroxy-1,4-benzoxazin-3-one-glucoside (DIBOA-Glc), the most abundant benzoxazinoid (BX) in cereal crops and the precursor of DIMBOA-Glc. BXs have dual functions in the rhizosphere, suppressing pathogens while recruiting PGPR.[48] In maize, DIMBOA-Glc is the main BX compound secreted by roots and was shown to stimulate the growth of certain rhizosphere bacteria in the Bacillaceae family.[49] DIBOA-Glc may have similar functions in wheat as DIMBOA-Glc in maize, as here we observed an enrichment of Bacillaceae in the biochar-treated rhizosphere (**Table S5**).

#### 2.4. Biochar-treated rhizosphere microbiome has shifted functions

##### *2.4.1. Shifted N and methane cycling*

PICRUSt2 predicted the enrichment of eight pathways in biochar-treated rhizosphere, including L-methionine salvage cycle I, nitrifier denitrification, biotin biosynthesis II, methanogenesis from acetate, methylaspartate cycle, glutaryl-CoA degradation, L-glutamate degradation V (via hydroxyglutarate), superpathway of mycolyl-arabinogalactan-peptidoglycan complex biosynthesis, and suppression of nine pathways, including polymyxin resistance, superpathway of glycol metabolism and degradation, coenzyme M biosynthesis I, superpathway of methylglyoxal degradation, glucose degradation (oxidative), 4-hydroxyacetophenone degradation, heparin degradation, spirilloxanthin and 2,2'-diketo-spirilloxanthin biosynthesis, coumarins biosynthesis (engineered) (**Fig 6A**).

Since biochar has been widely reported to mitigate $N_2O$ and $CH_4$ emissions in bulk soil, we paid close attention to N and $CH_4$ cycling genes in PICRUSt2 analysis, including those involved in N fixation, nitrification, assimilatory nitrate reduction, dissimilatory nitrate reduction, denitrification, glutamate synthesis, nitrate/nitrite transporter, and methane metabolism (**Fig S12-S13; Table S10**). Biochar increased nitrogen fixation genes (*nifD*, *nifH*, *nifK*) and nitrification genes (*amoA*, *amoB, amoC*, *hao*, *nxrA*, and *nxrB*), while decreasing denitrification genes (*nirK*, *nirS*, *norB*, *norC*, *nosZ*), although none was statistically significant (**Fig S12**). We also measured the abundance of *nifH*, *nxrA*, *nirK*, *nirS*, and *nosZ* using qPCR. qPCR confirmed PICRUSt2-predicted increases in *nifH* and *nxrA* and decrease in *nirK* ($p > 0.05$ for all) (**Fig 6B**). In contrast to PICRUSt2-predictions, qPCR identified increasing *nirS* and *nosZ* abundance in the biochar-treated rhizosphere ($p > 0.05$ for both). This discrepancy may reflect the relatively narrow scope of the used primers as our knowledge of N pathways in microorganisms continue to grow.

PICRUSt2 did not detect the *mcrA* gene encoding methyl-coenzyme M reductase complex subunit A in any sample. PICRUSt2 found insignificant increases in *mmoX* encoding soluble methane monooxygenase and *pmoA* encoding particulate methane monooxygenase subunit A in the biochar-treated rhizosphere (**Fig S13**). Furthermore, *pmoA* was more prevalent than *mmoX*. Consistently, qPCR did not detect *mcrA* in any sample, but found increased *pmoA* in the biochar-treated rhizosphere ($p > 0.05$) (**Fig 6B**). In addition, biochar may have influenced methylated substrates and thereby methyl-based methanogenesis,[50,51] as suggested by PICRUSt2-predicted enrichment (>1000 fold) of methionine salvage pathway (MSP) in the biochar-treated rhizosphere (**Fig 6A**). Coincidently, several MSP intermediates, S-adenosylmethioninamine, 5-Methylthioribose-1-phosphate (MTR-1-P) and 5-methylthioribulose-1-phosphate (MTRu-1-P), were detected in root exudates, while S-adenosylmethioninamine and a downstream product, 3-methylthiopropionic acid, were detected in the soil metabolome (**Table S3; Table S12**). These methylthio compounds were significantly associated with microbial cysteine and methionine metabolism in the biochar-treated rhizosphere (*p-adjusted* < 0.05) (**Table S9; Table S13**), and four PGPR showed high co-occurrence with 3-methylthiopropionic acid in the soil metabolome (*Roseomonas* > *Nevskia* > *Blastococcus* > *Bacillus*) (**Fig S14**).

##### *2.4.2. Broader changes in rhizosphere microbiome functions*

Biochar significantly affected 64 of the 506 exometabolites ($|Log2FC| > 1$; Mann-Whitney U test with Benjamini-Hochberg correction: $p < 0.05$) (**Fig 6C**). These 64 exometabolites fell into 14 ChemOnt classes, including 10 organooxygen compounds, 9 steroids and steroid derivatives, 8 fatty acyls, 7 prenol lipids, 4 tetrapyrroles and derivatives, 3 carboxylic acids and derivatives, 2 phenols, 2 keto acids and derivatives, 2 indoles and derivatives, 1 pyridines and derivatives, 1 pteridines and derivatives, 1 organic phosphoric acids and derivatives, 1 furofurans, 1 dihydrofurans, and 12 unclassed chemicals. Fifty-six of the 64 exometabolites were downregulated and eight were upregulated.

Among the 506 exometabolites, 180 compounds were also detected in root exudates collected by hydroponic secretion while 326 compounds were only found in the soil metabolome (**Fig 7A; Table S12**). It should be noted that the 326 compounds could be microbial metabolites or produced by plants in contact with rhizosphere microbes. The 326 soil exometabolites were mapped to 72 predicted microbial pathways (**Table S13**), and 27 of the 72 pathways were also observed when inferring microbe-root exudate interactions (**Table S9**). These 27 pathways fell into Central Carbon & Energy Metabolism (Glycolysis / Gluconeogenesis, Citrate cycle (TCA cycle), Pyruvate metabolism, Pentose phosphate pathway, Glyoxylate and dicarboxylate metabolism, Propanoate metabolism), Carbohydrate Metabolism (Fructose and mannose metabolism, Galactose metabolism, Starch and sucrose metabolism), Amino Acid Metabolism (Phenylalanine, tyrosine and tryptophan biosynthesis, Arginine and proline metabolism, Cysteine and methionine metabolism), Metabolism of Cofactors & Vitamins (Porphyrin and chlorophyll metabolism, Folate biosynthesis, Pantothenate and CoA biosynthesis, Biotin metabolism), Metabolism of Other Small Molecules (Pyrimidine metabolism, Glutathione metabolism), Membrane Transport (ABC transporters), and Cellular Processes & Stress Responses (Ferroptosis), among others.

After mapping the 326 soil exometabolites to 72 predicted microbial pathways (**Table S13**) and comparing microbe-root exudate co-occurrence with microbe-exometabolite co-occurrence (**Table S9**), we found biochar likely influenced microbially mediated biosynthesis of several vitamins, including biotin, phylloquinone, and menaquinone. Biotin, its immediate precursor dethiobiotin, and its activated intermediate biotinyl-5'-AMP were only detected in the soil metabolome (**Table S12)**. Coincidently, PICRUSt2 predicted that biochar promoted biotin biosynthesis II (**Fig 6A**) in the rhizosphere microbiome, and biotin showed very high co-occurrence probability with several PGPR, including *Brevundimonas* with biotin biosynthesis II pathway (**Fig S14**)[41]. Phylloquinone is a vitamin (K1) produced by plants and cyanobacteria. Phylloquinone and its direct percussor demethylphylloquinone were detected in both root exudates and the soil metabolome, but their reduced forms, demethylphylloquinol and phylloquinol, were only detected in the soil metabolome (**Table S3; Table S12**). Biochar significantly reduced demethylphylloquinone and phylloquinol in rhizosphere soils (**Fig 6C**). Consistently, PICRUSt2 predicted a reduction of NQO1 (K00355)[52] encoding NAD(P)H dehydrogenase (phylloquinone to phylloquinol) under the biochar treatment. These results likely reflect that biochar stimulated certain Cyanobacteria (discussed in **SI Section 2.3.1**) which use the photosystem I complex with phylloquinone as electron carrier for photosynthesis. Menaquinone is a vitamin (K12) produced by bacteria and some archaea. The direct precursor of menaquinone, 2-succinyl-5-enolpyruvyl-6-hydroxy-3-cyclohexene-1-carboxylic acid, was only detected in root exudates. In the menaquinone biosynthesis pathway, 2-succinyl-5-enolpyruvyl-6-hydroxy-3-cyclohexene-1-carboxylic acid is first converted to (1R,6R)-6-hydroxy-2-succinylcyclohexa-2,4-diene-1-carboxylate and then to 2-succinylbenzoate and finally to menaquinol. The two intermediates of menaquinone biosynthesis were only detected in the soil metabolome (**Table S12**), and biochar significantly reduced their abundance in the rhizosphere (**Fig 6C**). Coincidentally, PICRUSt2 predicted that biochar increased all but one genes involved in menaquinone biosynthesis, including MenH (K08680)[53] and MenC (K02549)[54] that catalyze the transformation of 2-succinyl-5-enolpyruvyl-6-hydroxy-3-cyclohexene-1-carboxylic acid to (1R,6R)-6-hydroxy-2-succinylcyclohexa-2,4-diene-1-carboxylate and 2-succinylbenzoate (**Fig S15**). Menaquinol, the bioactive reduced form of menaquinone, was detected in the soil metabolome (**Fig S15**).

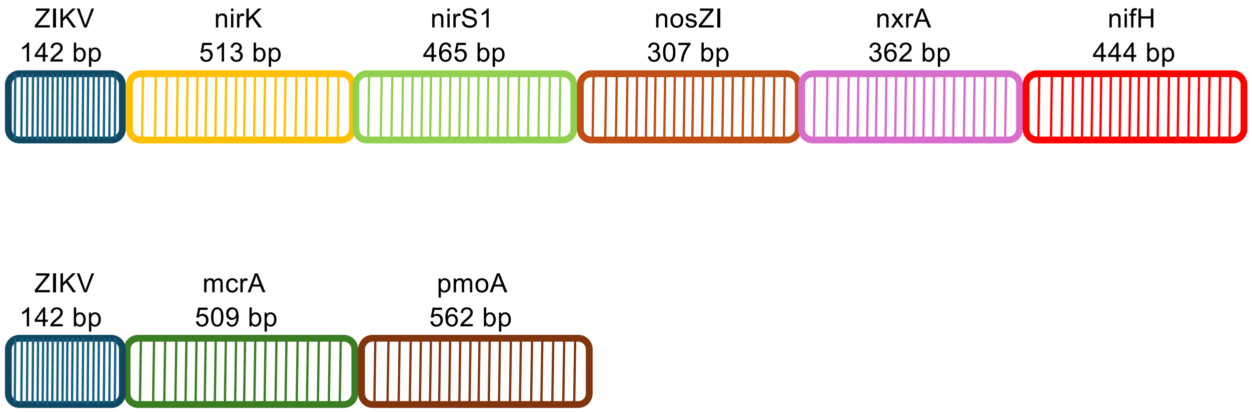


**Fig S1.** Schematic of positive controls (gBlocks, Integrated DNA Technologies, Inc.) of N-cycling (upper) and methane-cycling (bottom) genes used in qPCR. Gene fragments are shown as colored blocks with PCR amplicon sizes denoted.

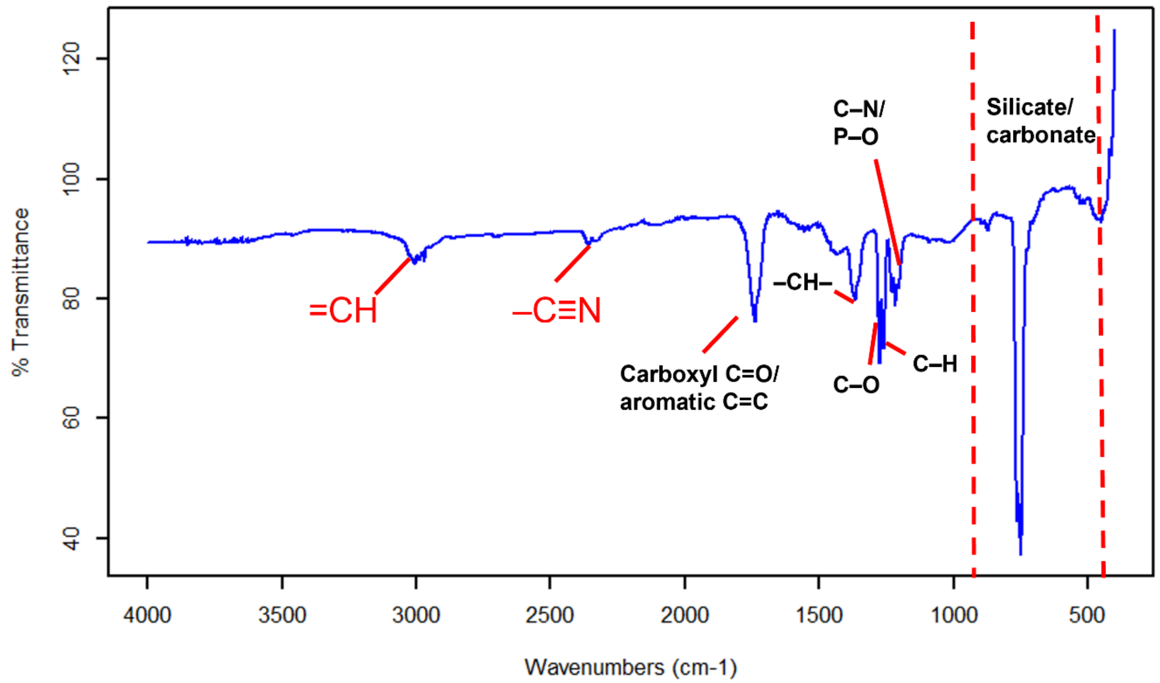


422

423 **Fig S2.** Full-range FTIR spectrum (400–4000 $cm^{-1}$) of the wheat biochar used in this study.

424

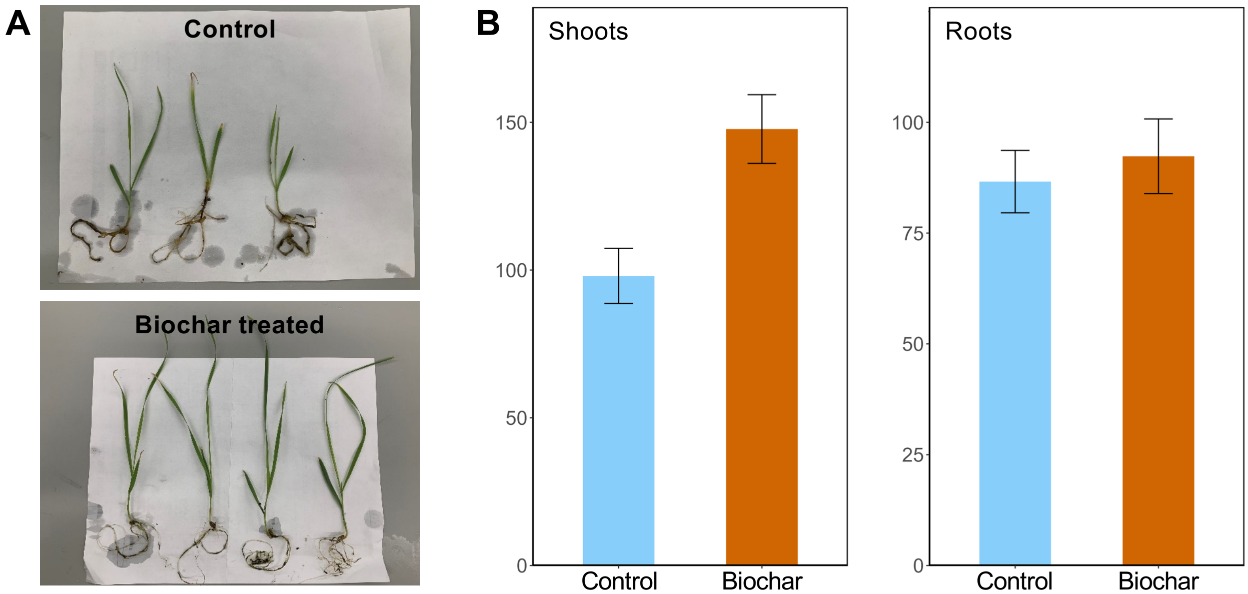


**Fig S3.** (A) Representative wheat plants grown with or without 0.25% wheat biochar in the EcoFAB experiment. (B) Fresh weight (mg) of shoots (left) and roots (right) of wheat plants.

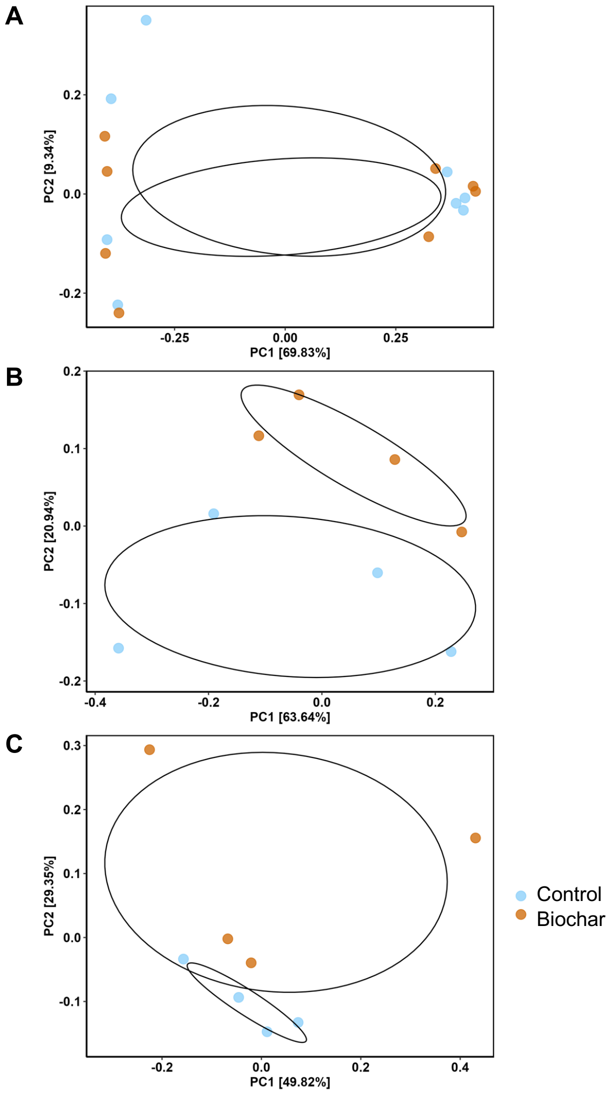


**Fig S4**. PCoA of Jaccard distance of root exudates detected in the control and biochar-treated plants after 21 days. The ellipses represent 95% confidence. (A) Data from two rounds of EcoFAB experiments separated primarily by the experiment round along the first coordinate, which explains 69.83% of the total variance (PERMANOVA: $p$ = 0.001 for experiment round; $p$ = 0.211 for treatment; $p$ = 0.145 for round x treatment). (B-C) Data from the (B) first-round and (C) second-round experiment was consistent and separated primarily by treatment on the second coordinate, which explains 20.94% and 29.35% of the total variance, respectively (PERMANOVA: $p$ = 0.221 and $p$ = 0.154 for treatment, respectively).

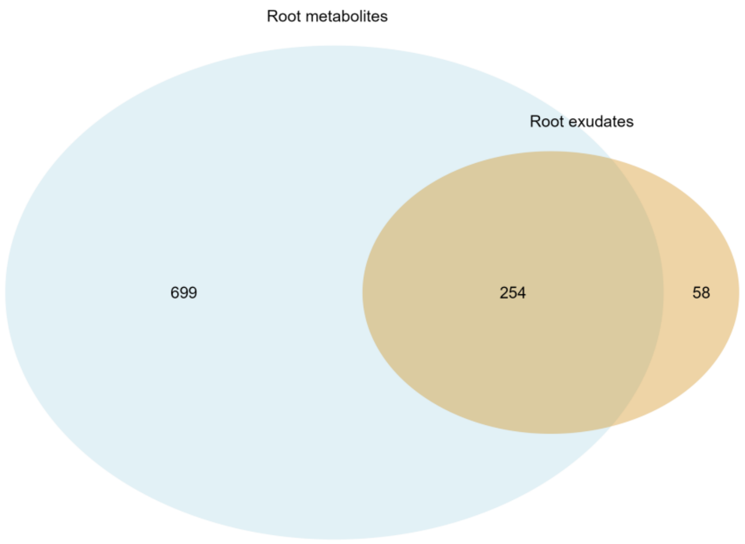


**Fig S5.** Venn diagram showing root endometabolites and root exudates of wheat plants grown under 0.25% wheat biochar. Root endometabolites were detected in a prior pot experiment with ESI Q-TOF HRMS[1], while root exudates were detected here in an EcoFAB experiment with Q Exactive HF Orbitrap HRMS.

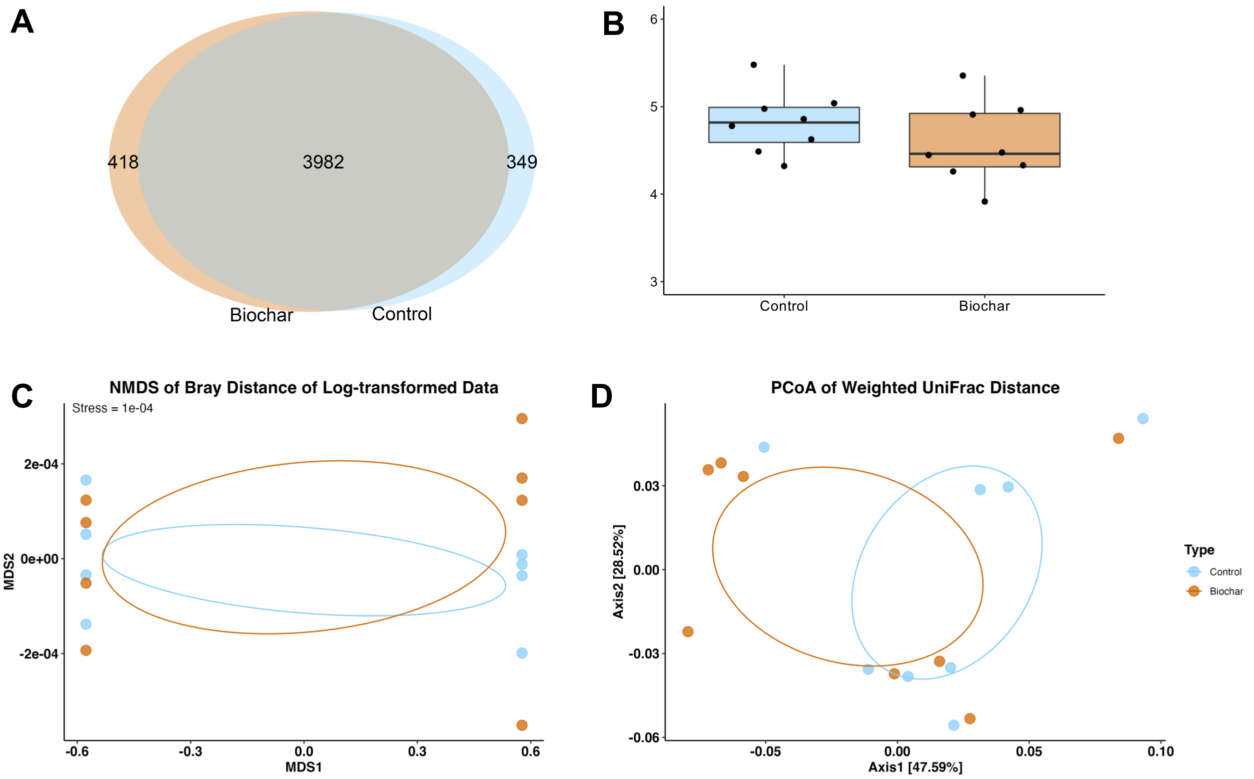


**Fig S6.** (A-B) Biochar slightly altered the alpha diversity of the rhizosphere microbiome, as shown by (A) the Venn diagram of microbial ASVs (i.e., richness) and (B) the Box-and-whisker plot of Shannon diversity. (C-D) Biochar significantly altered the beta diversity of the rhizosphere microbiome, evident in (C) the nonmetric multidimensional scaling (NMDS) analysis of Bray-Curtis distance of log-transformed ASV abundance (stress = 0.0001) and (D) the principal coordinates analysis (PCoA) of weighted UniFrac distance of ASV abundance (PERMANOVA test: $p < 0.001$ for experiment round; $p = 0.217$ for treatment; $p = 0.001$ for round $\times$ treatment).

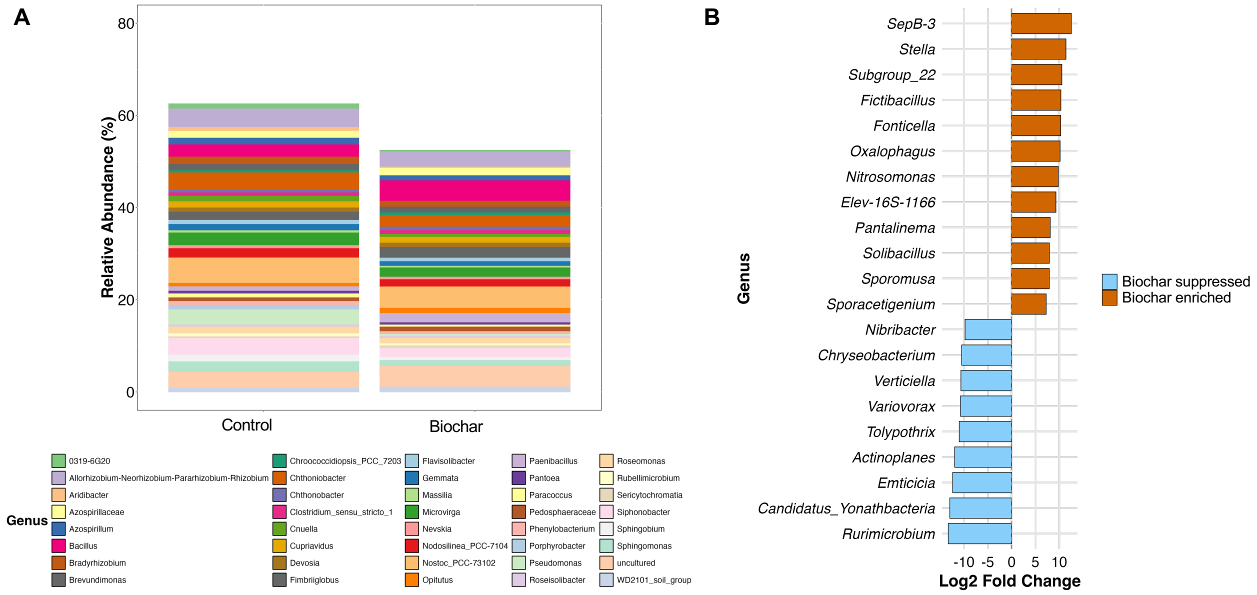


**Fig S7.** (A) Relative abundance of the top 40 rhizosphere microbial genera of the control and biochar-treated wheat plants. Data presents averages across biological replicates. (B) Microbial genera significantly influenced by 0.25% wheat biochar (|Log2FC| > 1, *p-adjusted* < 0.05).

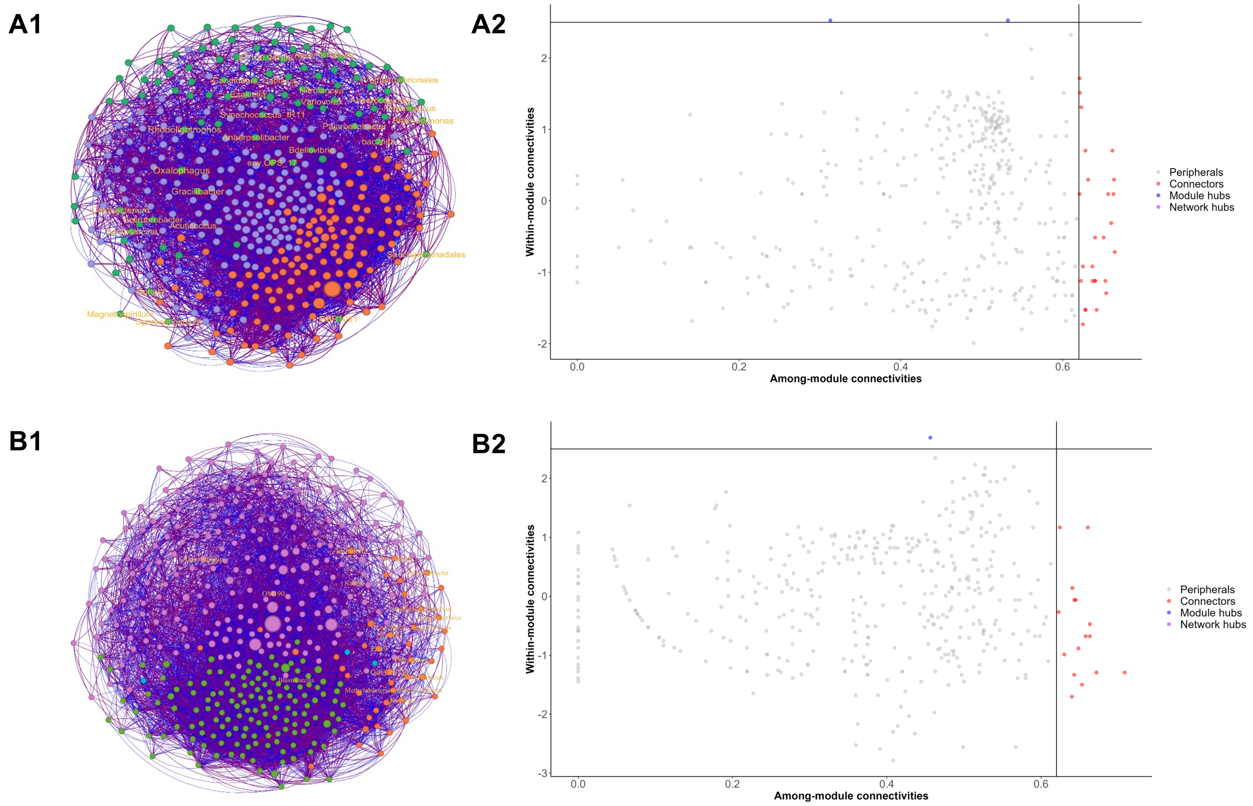


**Fig S8.** Network of microbial genera in the control rhizosphere (A1) and the biochar-treated rhizosphere (B1). Co-occurrence network was constructed using SparCC. Only significant correlations ($p$ < 0.05 from 1000 bootstrap resamples) were retained. Node colors indicate different modules. Node sizes are proportional to the betweenness values. Purple and blue edges indicate positive and negative correlations, respectively. Within-module (Zi) and between-module (Pi) connectivity of network nodes were calculated to identify network connectors and module hubs, i.e., keystone taxa, in the control rhizosphere (A2) and the biochar-treated rhizosphere (B2). Identified keystone taxa were labeled with genus identities in the networks (A1, B1).

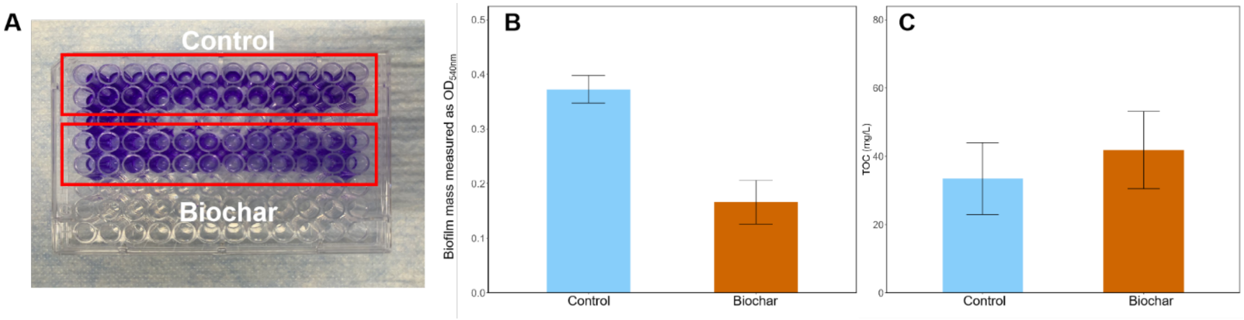


**Fig S9.** (A) Biofilm assay using *P. putida KT2440* in a 96-well microtiter plate. Wells in the upper two rows and the lower two rows contained root exudate cocktails collected from control and biochar-treated wheat plants, respectively. (B) Mass (in $OD_{540nm}$) of biofilm formed in the microtiter plate after 48 hours of static incubation at 27 °C. Data represents average and one standard deviation from 4 biological replicates ($p$ = 0.004). (C) DOC levels of control and biochar-induced root exudate cocktails were similar ($p$ = 0.607).

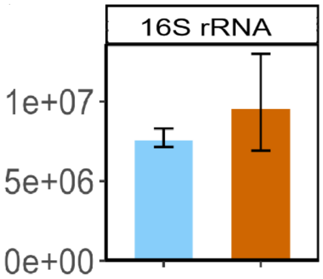


**Fig S10.** Abundance (copies/ng DNA) of eubacterial 16S rRNA gene in control (blue) and biochar-induced (brown) rhizosphere.

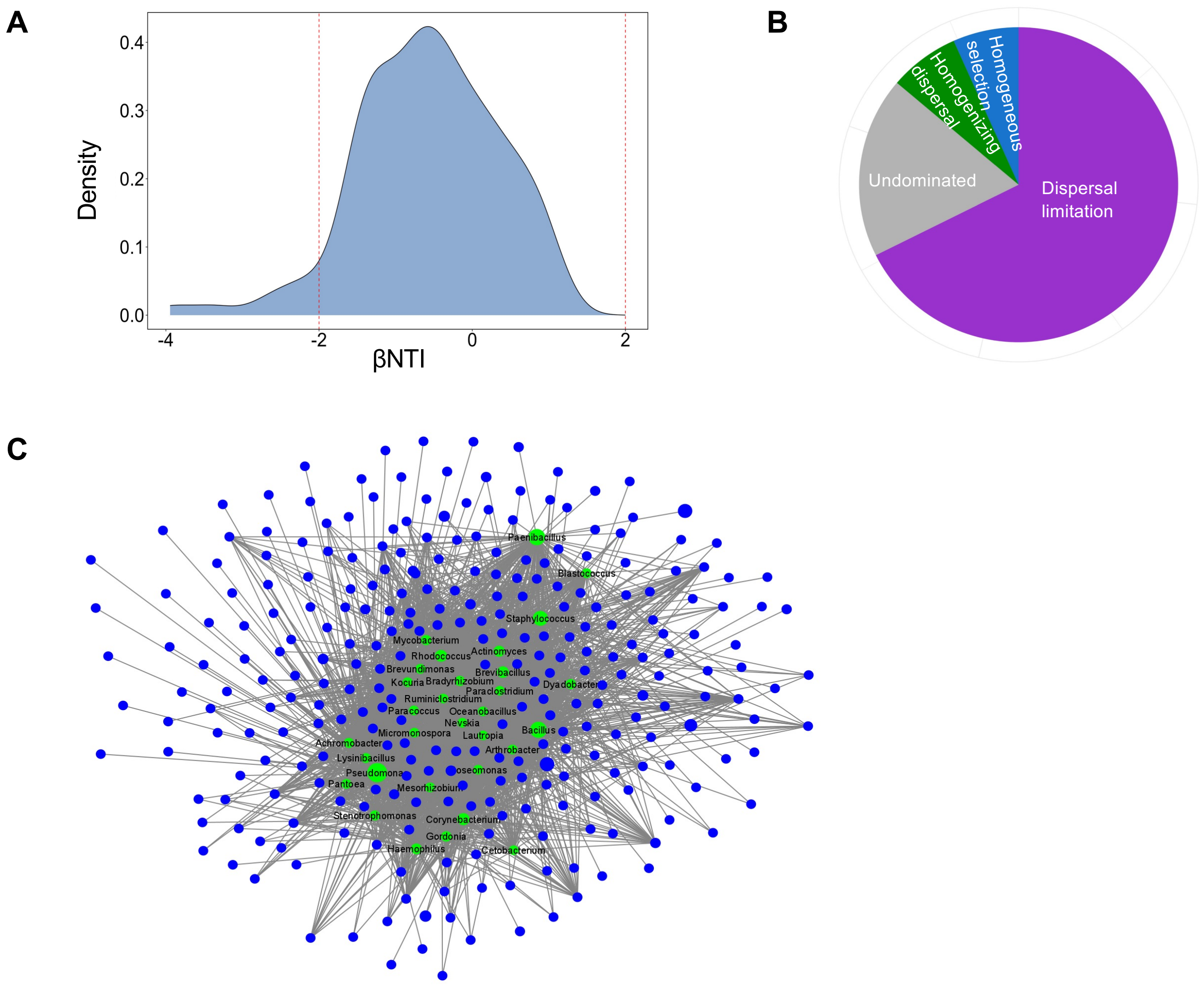


**Fig S11.** (A) Stochastic processes dominated the assembly of the biochar-rewired rhizosphere microbiome. (B) Pie chart of assembly processes with slice area proportional to their relative importance: dispersal limitation (stochastic, 63.11%) >> undominated (neither deterministic nor stochastic, 17.33%) >> homogenizing dispersal (stochastic, 6.67%) > homogenous selection (deterministic, 6.22%). Assembly mechanism and importance were estimated using βNTI and $RC_{BC}$. βNTI<−2: homogenous selection; βNTI>2: heterogenous selection; |βNTI|<2 and $RC_{BC}$<-0.95: homogenizing dispersal; |βNTI|<2 and $RC_{BC}$>0.95: dispersal limitation; |βNTI|<2 and |$RC_{BC}$|<0.95: undominated. (C) Associations between exudate compounds (blue) and microbial genera (green) inferred using co-occurrence network (OmicsNet 2.0). Node size is proportional to the betweenness value of a node.

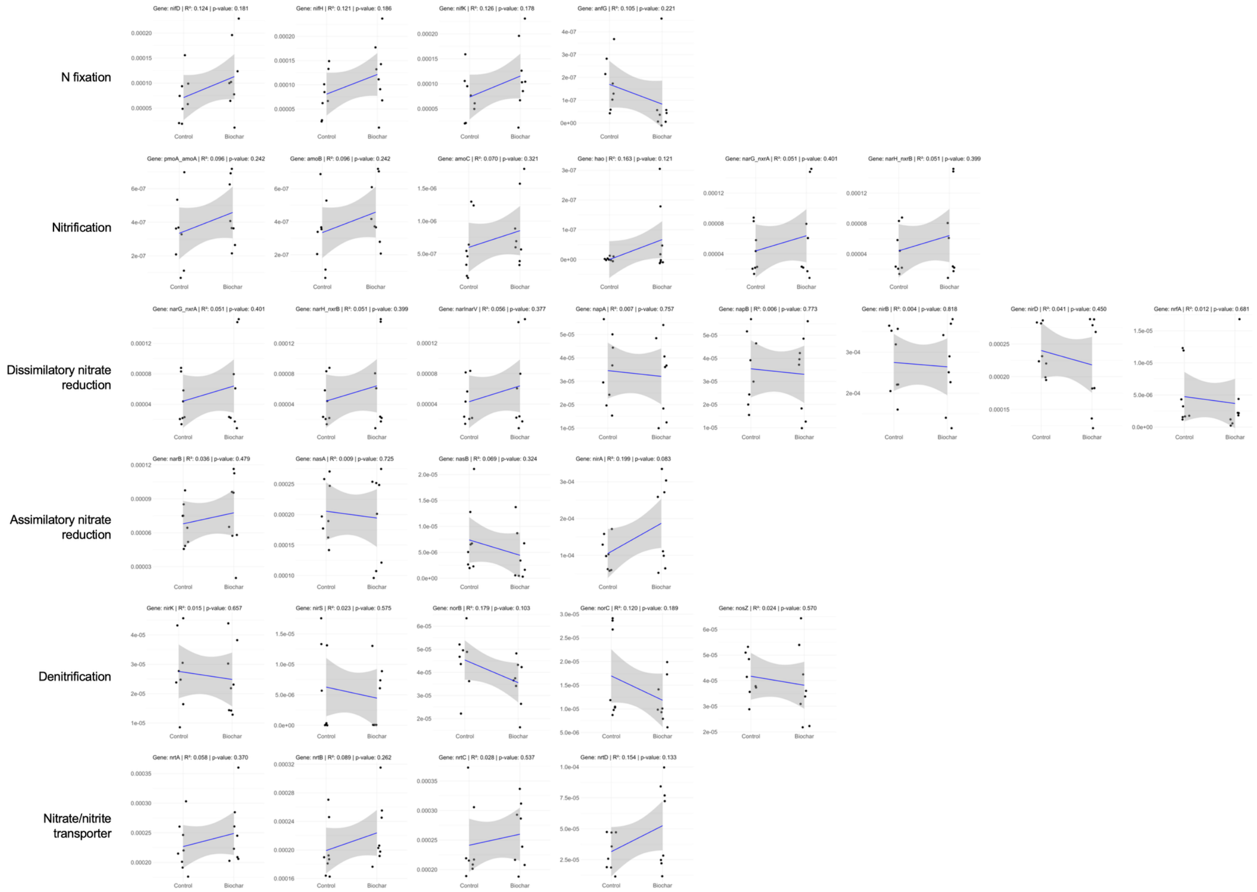


**Fig S12**. PICRUSt2 predicted relative abundance of N-cycling genes. Genes are grouped based on major pathways. For each subplot, linear regression $R^2$ and *p*-value are shown on the top, and the grey zone indicates the 95% confidence interval.

Methane oxidation

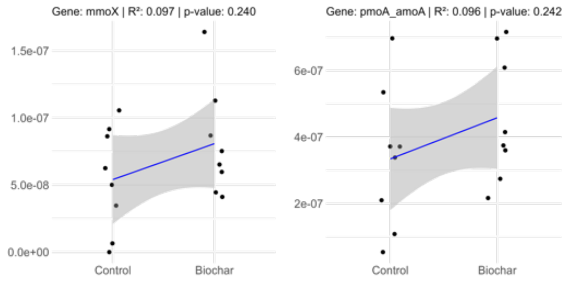


**Fig S13**. PICRUSt2 predicted relative abundance of methane-cycling genes. Genes are grouped based on major pathways (methanogenesis pathway not detected). For each subplot, linear regression $R^2$ and *p*-value are shown on the top, and the grey zone indicates the 95% confidence interval.

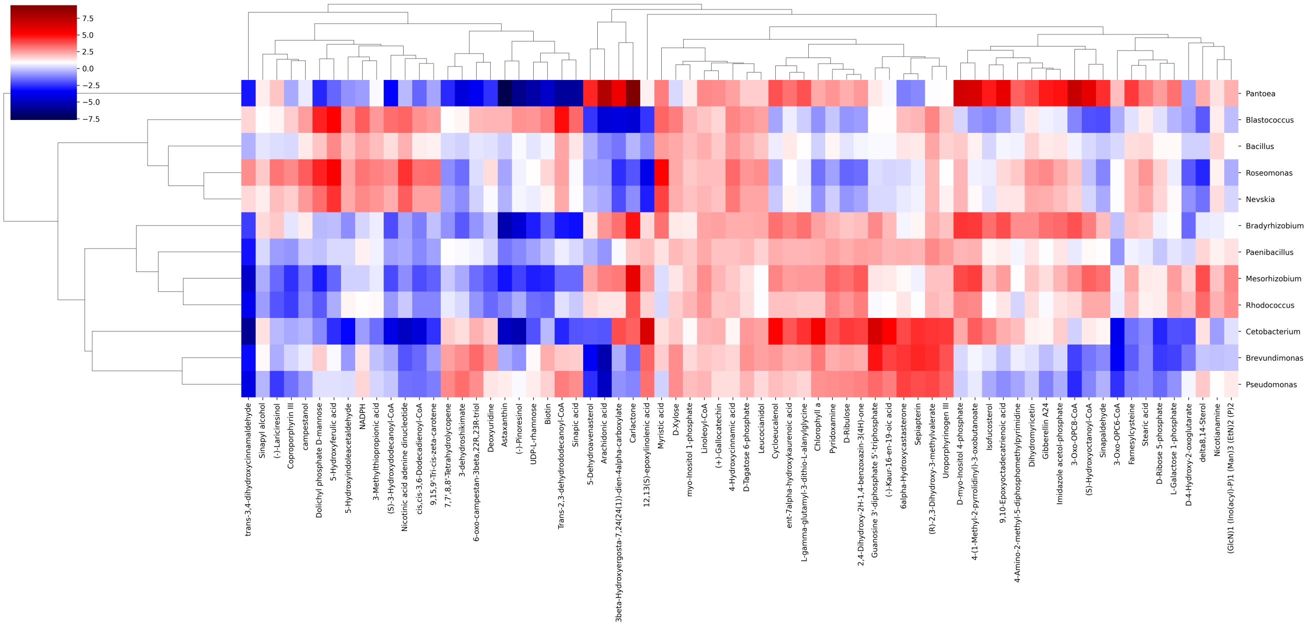


**Fig S14.** Associations between soil exometabolites (column) and bacterial genera (row) inferred by the mmvec neural network. Colors indicate the estimated log conditional probability (i.e., association strength) of observing a soil exometabolite compound given the presence of a microbial genus. Only those genera identified by both the OmicsNet 2.0 and mmvec approaches are shown here to ensure true positivity. Soil exometabolites are clustered based on their associations on microbial genera.

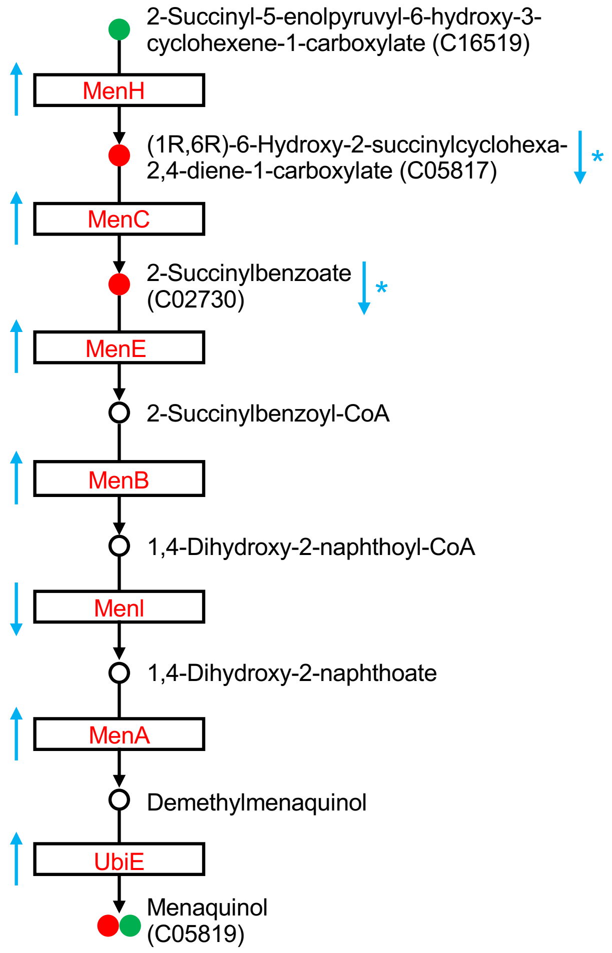


**Fig S15.** Integrating multi-omics data suggests upregulated menaquinol biosynthesis in the biochar-treated rhizosphere microbiome. Dots indicate compounds: Green, detected in root exudates; red, detected in the soil metabolome; open, undetected compounds. KEGG C numbers are listed for detected compounds. Boxes indicate microbial enzymes inferred by PICRUSt2: Red, regulated by biochar; black, no change after the biochar treatment. Blue directed arrows indicate up- or down-regulation of enzymes or compounds, with asterisks representing significant changes (also see **Fig 6C**).

(12) Bolyen, E.; Rideout, J. R.; Dillon, M. R.; Bokulich, N. A.; Abnet, C. C.; Al-Ghalith, G. A.; Alexander, H.; Alm, E. J.; Arumugam, M.; Asnicar, F.; Bai, Y.; Bisanz, J. E.; Bittinger, K.; Brejnrod, A.; Brislawn, C. J.; Brown, C. T.; Callahan, B. J.; guez, A. x000E9 s M. C.-R. x000ED; Chase, J.; Cope, E. K.; Silva, R.; Diener, C.; Dorrestein, P. C.; Douglas, G. M.;

Durall, D. M.; Duvallet, C.; Edwardson, C. F.; Ernst, M.; Estaki, M.; Fouquier, J.; Gauglitz, J. M.; Gibbons, S. M.; Gibson, D. L.; Gonzalez, A.; Gorlick, K.; Guo, J.; Hillmann, B.; Holmes, S.; Holste, H.; Huttenhower, C.; Huttley, G. A.; Janssen, S.; Jarmusch, A. K.; Jiang, L.; Kaehler, B. D.; Bin Kang, K.; Keefe, C. R.; Keim, P.; Kelley, S. T.; Knights, D.; Koester, I.; Kosciolek, T.; Kreps, J.; Langille, M. G. I.; Lee, J.; Ley, R.; Liu, Y.-X.; Loftfield, E.; Lozupone, C.; Maher, M.; Marotz, C.; Martin, B. D.; McDonald, D.; McIver, L. J.; Melnik, A. V.; Metcalf, J. L.; Morgan, S. C.; Morton, J. T.; Naimey, A. T.; Navas-Molina, J. A.; Nothias, L. F.; Orchanian, S. B.; Pearson, T.; Peoples, S. L.; Petras, D.; Preuss, M. L.; Pruesse, E.; Rasmussen, L. B.; Rivers, A.; Robeson, M. S.; Rosenthal, P.; Segata, N.; Shaffer, M.; Shiffer, A.; Sinha, R.; Song, S. J.; Spear, J. R.; Swafford, A. D.; Thompson, L. R.; Torres, P. J.; Trinh, P.; Tripathi, A.; Turnbaugh, P. J.; Ul-Hasan, S.; Hooft, J. J. J.; Vargas, F.; zquez-Baeza, Y. V. x000E1; Vogtmann, E.; von Hippel, M.; Walters, W.; Wan, Y.; Wang, M.; Warren, J.; Weber, K. C.; Williamson, C. H. D.; Willis, A. D.; Xu, Z. Z.; Zaneveld, J. R.; Zhang, Y.; Zhu, Q.; Knight, R.; Caporaso, J. G. Reproducible, Interactive, Scalable and Extensible Microbiome Data Science Using QIIME 2. *Nature Biotechnology* **2019**, 1–6. https://doi.org/10.1038/s41587-019-0209-9.

(21) Lewin, S.; Francioli, D.; Ulrich, A.; Kolb, S. Crop Host Signatures Reflected by Co-Association Patterns of Keystone Bacteria in the Rhizosphere Microbiota. *Environmental*

*Microbiome* **2021**, *16* (1), 18. https://doi.org/10.1186/s40793-021-00387-w.
(22) Banerjee, S.; Schlaeppi, K.; Heijden, M. G. A. Keystone Taxa as Drivers of Microbiome Structure and Functioning. *Nat Rev Micro* **2018**, 1–10. https://doi.org/10.1038/s41579-018-0024-1.
(23) Layeghifard, M.; Hwang, D. M.; Guttman, D. S. Disentangling Interactions in the Microbiome: A Network Perspective. *Trends in Microbiology* **2017**, *25* (3), 217–228. https://doi.org/10.1016/j.tim.2016.11.008.
(24) Bustin, S. A.; Benes, V.; Garson, J. A.; Hellemans, J.; Huggett, J.; Kubista, M.; Mueller, R.; Nolan, T.; Pfaffl, M. W.; Shipley, G. L.; Vandesompele, J.; Wittwer, C. T. The MIQE Guidelines: Minimum Information for Publication of Quantitative Real-Time PCR Experiments. *Clinical Chemistry* **2009**, *55* (4), 611–622. https://doi.org/10.1373/clinchem.2008.112797.
(25) Gillespie, C. J.; Sit, T. L.; Loucanides, A. J.; Hu, J.; Matusko, B. E.; Terwilliger, N. R.; Shi, W.; Steffen, S. M.; Woodley, A. L.; Hu, S. Multi-Amplicon Nitrogen Cycling Gene Standard: An Innovative Approach for Quantifying N-Transforming Soil Microbes in Terrestrial Ecosystems. *Soil Biology and Biochemistry* **2024**, *195*, 109461. https://doi.org/10.1016/j.soilbio.2024.109461.
(26) You, Y.; Hilpert, M.; Ward, M. J. Detection of a Common and Persistent Tet(L)-Carrying Plasmid in Chicken-Waste-Impacted Farm Soil. *Appl. Environ. Microbiol.* **2012**, *78* (9), 3203–3213. https://doi.org/10.1128/AEM.07763-11.
(27) Suzuki, M. T.; Taylor, L. T.; Delong, E. F. Quantitative Analysis of Small-Subunit rRNA Genes in Mixed Microbial Populations via 5′-Nuclease Assays. *Applied and Environmental Microbiology* **2000**.
(28) Burmølle, M.; Hansen, L. H.; Sørensen, S. J. Establishment and Early Succession of a Multispecies Biofilm Composed of Soil Bacteria. *Microb Ecol* **2007**, *54* (2), 352–362. https://doi.org/10.1007/s00248-007-9222-5.
(29) Rodrigues, D. F.; Elimelech, M. Toxic Effects of Single-Walled Carbon Nanotubes in the Development of E. Coli Biofilm. *Environ. Sci. Technol.* **2010**, *44* (12), 4583–4589. https://doi.org/10.1021/es1005785.
(30) Magnúsdóttir, S.; Heinken, A.; Kutt, L.; Ravcheev, D. A.; Bauer, E.; Noronha, A.; Greenhalgh, K.; Jäger, C.; Baginska, J.; Wilmes, P.; Fleming, R. M. T.; Thiele, I. Generation of Genome-Scale Metabolic Reconstructions for 773 Members of the Human Gut Microbiota. *Nat Biotechnol* **2017**, *35* (1), 81–89. https://doi.org/10.1038/nbt.3703.
(31) Zhou, G.; Pang, Z.; Lu, Y.; Ewald, J.; Xia, J. OmicsNet 2.0: A Web-Based Platform for Multi-Omics Integration and Network Visual Analytics. *Nucleic Acids Research* **2022**, *50* (W1), W527–W533. https://doi.org/10.1093/nar/gkac376.
(32) Morton, J. T.; Aksenov, A. A.; Nothias, L. F.; Foulds, J. R.; Quinn, R. A.; Badri, M. H.; Swenson, T. L.; Goethem, M. W.; Northen, T. R.; Vázquez-Baeza, Y.; Wang, M.; Bokulich, N. A.; Watters, A.; Song, S. J.; Bonneau, R.; Dorrestein, P. C.; Knight, R. Learning Representations of Microbe–Metabolite Interactions. *Nat Chem Biol* **2019**, 1–12. https://doi.org/10.1038/s41592-019-0616-3.

2920.12149.
